# Dimeric Alix nucleates ESCRT-III CHMP4 polymerization

**DOI:** 10.64898/2026.05.22.727116

**Authors:** Nolwenn Miguet, Carlos Contreras-Martel, Sourav Maity, Els Pardon, Stéphane Frémont, Pauline Macheboeuf, Christine Chatellard, Jean-Marie Bourhis, Jean-Philippe Kleman, Andréa Dessen, Remy Sadoul, Christophe Chipot, Jan Steyaert, Wouter H Roos, Arnaud Echard, Winfried Weissenhorn

## Abstract

Alix is a key adaptor protein of the endosomal sorting complex required for transport (ESCRT) membrane remodeling machinery. Although Alix-mediated recruitment of the ESCRT-III subunit CHMP4 is well established, the molecular mechanisms underlying Alix activation and ESCRT-III polymerization remain poorly understood. Here, we present the crystal structure of the dimeric Alix V-domain (Alix-V) in complex with a llama nanobody. Dimerization is mediated by domain swapping, generating an X-shaped flexible conformation. We demonstrate that Alix forms dimers *in vivo* and provide evidence for the recruitment of dimeric Alix to plasma membrane repair sites and the cytokinetic midbody. Notably, a mutation disrupting Alix dimerization impairs plasma membrane repair. Furthermore, high-speed AFM experiments reveal that dimeric Alix, but not the monomeric form, nucleates CHMP4B filament polymerization. Our data establish that dimeric Alix is the active form responsible for nucleating two ESCRT-III CHMP4 filaments, whose geometry creates a platform for the recruitment of downstream ESCRT-III components required to assemble the active membrane remodeling and fission machinery.

## Introduction

The ALG-2-interacting protein X (Alix) ^1^ is an important adaptor protein of the endosomal sorting complex required for transport (ESCRT) machinery, which catalyzes many topologically similar membrane remodeling processes that terminate by membrane fission ^2–7^.

The ESCRT machinery is composed of five complexes, ESCRT-0, ESCRT-I, ESCRT-II, ESCRT-III and VPS4, of which ESCRT-III and VPS4 are universal to all ESCRT-catalyzed processes ^2^. Humans express eight ESCRT-III proteins named CHMP1 to 8 that can comprise several isoforms per member, including three CHMP4 paralogs ^5^. CHMP recruitment to membranes is thought to activate and open their closed conformation ^8–11^, which in turn leads to polymerization if CHMP1B ^12^, CHMP4 (Snf7) ^13–15^ and CHMP2A-CHMP3 ^16^ into loose or helical filaments. Notably, CHMP4 polymerization is central to all known ESCRT-catalyzed processes serving as a platform to recruit downstream ESCRT-III CHMP2 (Vps2) and CHMP3 (Vps24) ^17, 18,19^, which in turn were suggested to cap CHMP4 (Snf7) polymerization ^20^.

Although it is yet unknown what triggers the conformational change from the closed to the open ESCRT-III conformation, it is clear that ESCRT-III can be recruited by ESCRT-II ^21^, which forms a Y-shaped complex with two VPS25/EAP20 protomers ^22–24^ that interact with CHMP6 (Vps20) ^25^, which then recruits CHMP4 (Snf7) ^17^. The second ESCRT-III recruiter, Alix (Bro1) can directly engage CHMP4 (Snf7) without requiring ESCRT-II and CHMP6 (Vps20) ^26–29^. Alix, is composed of three domains, an N-terminal Bro1-domain, a central V-domain and a proline-rich domain (PRD) ^30^. The Bro1 domain binds to the C-termini of all CHMP4 paralogs ^31^, while the V-domain interacts with viral late domains ^32,33^, ubiquitin ^34^ and the syntenin/syndecan-4 complex ^35^. The PRD contains multiple interaction sites for diverse adaptors including ESCRT-I ^36,37^ and ALG-2 ^38^, the latter recruiting Alix to membranes in a Ca^2+^-dependent manner ^39^. The PRD was further suggested to keep Alix in an auto inhibited, inactive state in the cytosol by folding back onto the Bro1 and V domains thereby inhibiting viral late domain binding ^40^. Auto inhibition was proposed to be released via phosphorylation of PRD ^41^. Alternatively, Alix may be maintained in an autoinhibited state through hyperphosphorylation of its proline-rich domain (PRD) ^42^, which in turn promotes the formation of biomolecular condensates enriched in Alix and CHMP4 ^43^. Moreover, activation of Alix has been proposed to involve dimerization via its V-domain, generating an elongated conformation that can bridge two CHMP4B filaments *in vitro* ^44^. Despite the potential Alix modifications and interactions, it is yet unclear which of these regulatory mechanisms activate the closed conformation of Alix, which we hypothesize to lead to Alix dimerization.

Here we set out to determine the structure and function of Alix dimerization and its role in CHMP4 polymerization. We present the crystal structure of the dimeric Alix V-domain in complex with a llama nanobody. Dimerization is achieved via domain exchange generating an X-shaped dynamic structure. In agreement with a physiological role of dimeric Alix, we detected dimeric Alix in membrane fractions of HEK cells. Furthermore, we identified two llama nanobodies that bind to the Alix V-domain: NB89 is specific for inactive monomeric cytosolic Alix, while NB611 specifically detects the activated dimeric conformation of Alix present at membranes during ESCRT-catalyzed functions. Accordingly, we demonstrate that NB611 but not NB89 detects Alix recruitment to plasma membrane repair sites of HEK 293 cells and to the midbody of HeLa cells during cytokinesis, two processes that require ESCRT-III recruitment, activation and polymerization. Finally, using high-speed AFM we show that only dimeric Alix and not its monomeric form is able to nucleate CHMP4B polymerization *in vitro.* Together, our data establish that Alix recruits ESCRT-III in a similar way as ESCRT-II by coordinating two CHMP4 filaments. We propose that the geometries of the two CHMP4 filaments assembled on membranes is crucial for downstream ESCRT-III assembly and membrane remodeling.

## Results

### Structure of the dimeric Alix-V

Alix has been previously shown to dimerize via its V-domain ^30,44,45^. However, purified AlixΔPRD monomers do not dimerize in a concentration-dependent way, as confirmed by analytical ultracentrifugation (**Figure S1A**). Likewise, AlixΔPRD dimers do not dissociate into monomers at nanomolar concentrations (**Figure S1B**). This suggests that structural constraints must regulate Alix dimerization. In order to decipher Alix dimerization via its V-domain, we crystallized dimeric Alix-V (residues 358-714) containing two mutations of solvent-exposed residues, S568A and S577A, but these crystals yielded anisotropic diffraction data. To improve crystal diffraction, we generated Alix-V llama nanobodies by immunizing llamas with dimeric Alix-V. Although the isolated V-domain nanobodies were not specific for dimeric Alix-V and interacted as well with monomeric Alix-V, their interaction profiles with AlixΔPRD and full-length Alix differed (**Figure S2A**). We selected nanobody NB79 which interacted with AlixΔPRD (**Figure S2A and B**)(**Table S1**) and formed a stable complex with Alix-V as confirmed by size exclusion chromatography (SEC) and native gel analyses (**Figures S2C and D**). This procedure generated higher quality crystals. The structure of dimeric Alix-V-NB79 complexes was solved by molecular replacement and the model comprising residues (366-709) was refined to a resolution of 2.67 Å (**Table S2**). The structure exhibits an extended cross-shaped topology resembling a head-to-head configuration of two V protomers in complex with NB79 positioned at both ends of arms 1 and 1* **(Figure 1A**) being stabilized by a network of polar interactions (**Figure S2E**). Dimerization is achieved by domain exchange of the N-terminal helix 11 (α11) of arm 1, which interacts with arm 1* α18 and α19 of the second protomer. This shows that α18 and α19 of the monomeric V-domain need to detach from α11 and open into an extended structure in order to dimerize (**Figure 1B**). Both protomers are bridged by two connecting regions; notably hinge 1, which is formed by residues A533 to E547 connecting α17 to α18 and hinge 2, which is composed of residues Q641 to N643 connecting α19 and α20 (**Figure 1C**). This arrangement forms the two extended arms of the X-like structure (**Figure 1B**), which adopt an elongated (20.5 nm) flat structure as viewed from the side **(Figure 1D).** The structures of the individual V-domain arms are similar to the ones in the V-domain monomer structure (pdb 2OEX) and Cα atoms can be superimposed with an overall r.m.s.d. of 1.96 Å, although substantial structural flexibility is observed for the tips of arm 1 and 1* with the positions of α11, α 18 and α 19 deviating up to 14Å (E597) between monomeric Alix-V and the dimeric form (**Figures 1E and S3A**). This structural plasticity may in turn facilitate opening of the arm to drive dimerization.

**Figure 1.**
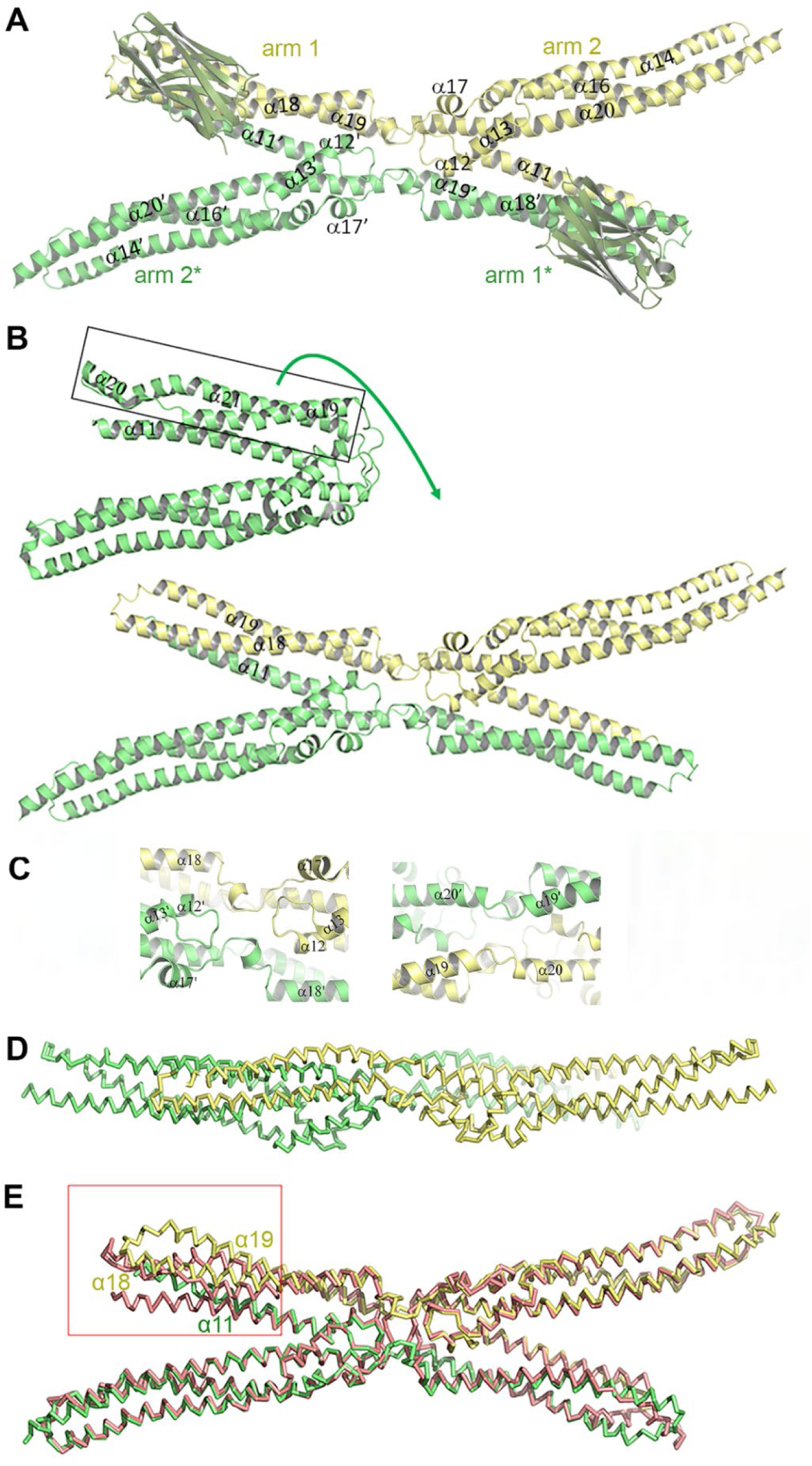
Structure of dimeric Alix-V. **(A)** Ribbon representation of dimeric Alix-V in complex with NB79. Secondary structure elements are labelled; the two protomers are shown in green and yellow. **(B)** Ribbon representation of monomeric (green; pdb 2OEX) and dimeric Alix-V, highlighting the conformational changes leading to Alix-V dimerization. Helices α29, α20 and α21 (corresponding to Alix-V dimer α18 and α19) must detach from α11 opening the V monomer conformation into an extended conformation in order to dimerize by domain exchange; movement indicated by green arrow. **(C)** Close-up of the hinge regions 1 and 2 (left and right panels) connecting α17 and α18 (hinge 1) and α19 to α20 (hinge 2) thereby joining the four arms of the X-like structure. **(D)** Cartoon representation of Alix-V rotated by 180° reveals a flat structure spanning the distance of 20 nm along the arm 2 axis. **(E)** Super positioning of the Cα atoms of V-domain monomers (left arms 1 and 2*, pdb 2OEV and right arms 2 and 1*, pdb 2OEX) demonstrates an excellent fit for arms 2 and 2* and part of arm 1 and 1*; however, the tips of arm 1 (circled in red) and 1* show substantial conformational plasticity with maximum displacements of helices by 14 Ǻ.

The mode of dimerization was confirmed by introducing pairs of cysteine mutations within the hinge region, i.e., N538C and A540C and V408C and A540C as well as cysteine pairs (V378C and N577C) crosslinking the N-terminal V-domain α11 of one protomer to α18 of the second protomer. Notably the Cα atoms of V378 and N577 are distanced by 11 and 12 Å in monomeric V structures (pdb 2OEX and 2OEV). All three sets of cysteine pairs led to the formation of disulfide-linked dimers as indicated by high molecular weight bands corresponding to dimeric forms detected by SDS PAGE analyses under non-reducing conditions (**Figures S3B and C**).

### Molecular dynamics of Alix-V

We next analyzed the flexibility of the X-shaped structure by performing three independent molecular dynamics (MD) simulations extending over 1 µs. Plotting the variability of the different angles α1 (arm 2 and arm 2*), α2 (arm 1 and arm 2; α2*, arm 1* and arm 2*) and α3 (arm 2 and arm 1*; α3*, arm 1 and arm 2*) revealed angles of 180°, 160° and 20°, respectively (**Figure S4A**). MD simulations showed α1 angle variations from 150° to 180° **(Figure S4B)**, α2 deviations between 120° and 180° with two runs stabilizing the conformational angle at 150° and one at 175° **(Figure S4C)**. The same flexibility could be observed for the symmetry-related angle α2* demonstrating fluctuations between 120° and 180°, one run stabilizing the conformation at ∼ 180° and two at ∼155° **(Figure S4D)**. α3 and its symmetry-related angle α3* showed further conformational variations leading to “closed” conformations with angles of ∼10°, and “open” ones with angles ranging from ∼40° to 55° (**Figures S4E, F**). Next, we used MD simulation on AlixΔPRD to analyze the capacity of α11 of the V domain to detach from α18 and α19, required for V-domain mediated dimerization. Plotting the RMSF from five different simulations showed that α11 exhibits moderate flexibility, with values ranging from 5 Å to <2 Å, which does not help to explain the dimerization-induced domain exchange. Only the loop connecting α14 to α15 revealed high flexibility with an RMSF up to 10 Å (**Figure S4G**). Thus, MD simulation analyses did not provide specific clues as to how domain exchange is achieved. However, it confirmed that the connecting hinges 1 and 2 allow for substantial flexibility of the X-shaped structure, which may be required for positioning diverse V-domain ligands such as viral late domains, syntenin and ubiquitin at ESCRT-coordinating membrane associated complexes.

### Membrane-associated Alix is dimeric and recognized by llama nanobody NB611

We co-expressed Flag-Alix and Myc-Alix to test their interaction in eukaryotic cells by co-IP, which were performed employing cytosolic and membrane fractions of HEK cells. IP of Flag-Alix allowed the detection of Myc-Alix in the membrane fraction but not in the cytosolic fraction (**Figure 2A**). Reciprocally, IP of Myc-Alix revealed Flag-Alix in the membrane fraction but not in the cytosolic fraction (**Figure 2B**). Together, this showed that Alix dimers were essentially present in membrane fractions. Although immunization with dimeric Alix-V did not produce nanobodies that can differentiate between monomers and dimers, we dissected binding of the obtained nanobodies to full-length Alix and monomeric AlixΔPRD (**Figure S2A**). Based on the ELISA assays we selected NB611 and NB89 due to their differential binding to AlixΔPRD and full-length Alix. While NB89 recognized both AlixΔPRD and full-length Alix, NB611 interacted only with AlixΔPRD suggesting that the presence of the PRD domain in full-length Alix prevents interaction (**Figure S2A**) consistent with full-length Alix in an inactive conformation. In line with two conformations of Alix, we show that NB611 immunoprecipitated Flag-Alix predominantly from the membrane fraction and not from the cytosolic fraction (**Figure 2C**), suggesting further that NB611 recognizes only membrane-associated active Alix and not inactive Alix that has been suggested to be monomeric and autoinhibited ^40^. We next analyzed full-length recombinant Alix by SAXS coupled to SEC-MALLS (**Figure S5A**) in order to test its conformational state. Guinier analysis of the scattering curve revealed a radius of gyration (Rg) of 46.4 Å consistent with SEC-MALLS (**Figures S5A, B and C**). In the dimensionless Kratky plot the curve reached a single well-defined peak around ∼1.3–1.4. for q.Rg values around 2.6 and showed a slow decay at q.Rg values above 4 (**Figure S5D**), in agreement with a mainly globular fold and some disordered regions, indicative of PRD dissociating from the Bro1-V domain in solution. PRD dissociation is further confirmed by the pair distance distribution function (PDDF) revealing a Dmax of 192 Å (**Figure S5E**) as well as by ab initio modelling with Dammin. The latter produced an ensemble of similar models (**Figure S5F**) with the best model fitting the corresponding experimental data with a discrepancy χ² of 0.73 and revealing a V-shaped envelope with extensions at both arms (**Figure S5G**). Notably, this conformation is incompatible with the proposed monomeric closed, autoinhibited Alix conformation with PRD folding over the V and Bro1 domains ^41^ (**Figure S5H**). We conclude that full-length recombinant Alix is a mixture of open and closed conformations *in vitro,* which renders the unambiguous determination of further biophysical characterization of NB611 binding *in vitro* inconclusive. However, in order to use NB611 and NB89 in cellular assays, we determined that both nanobodies have similar nanomolar binding K_D_s for monomeric and dimeric AlixΔPRD (**Table S1 and Figure S2B**) indicating that both nanobodies can be employed in comparative cellular assays.

**Figure 2.**
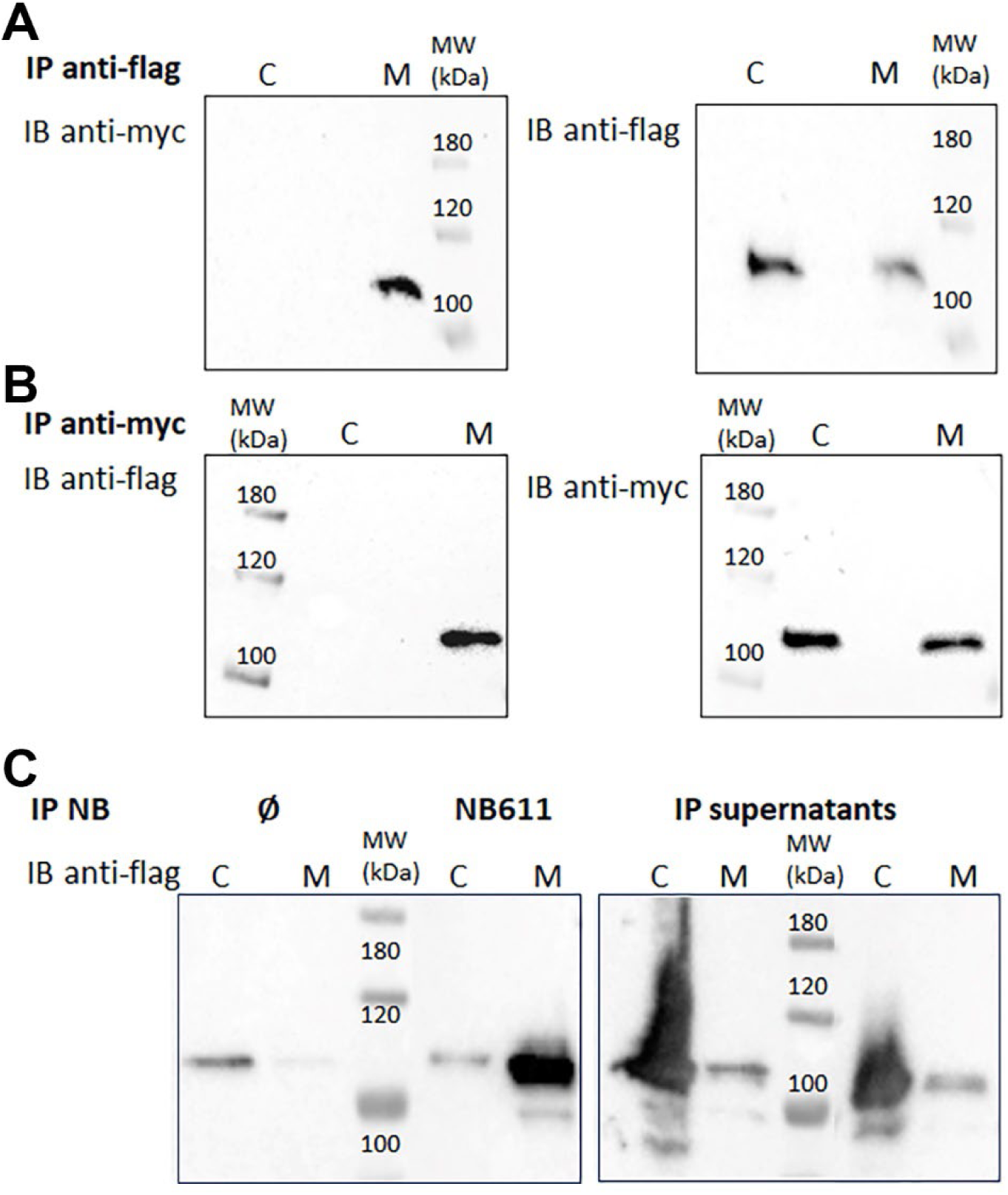
Membrane-associated but not cytosolic Alix is dimeric and recognized by llama nanobody NB611. **(A)** HEK cells were transfected with plasmids encoding Flag-Alix and Myc-Alix and were fractioned into cytosolic (C) and membrane (M) fractions. (left panel) Western blots of immune precipitation with anti-Flag beads and detection of Alix with an anti-Myc antibody revealed the presence of dimeric Alix in the membrane, but not cytosolic fraction. (right panel) Western blot with anti-flag antibodies detects Flag-Alix in both cytosolic and membrane fractions. **(B)** (left panel) Same as in (A) but Western blots of immune precipitation with anti-Myc beads and detection of dimeric Alix with an anti-Flag antibody revealed the presence of Alix in the membrane, but not cytosolic fraction. (right panel) Western blot with anti-Myc antibodies stains Myc-Alix in both cytosolic and membrane fractions, as expected. **(C)** Flag-Alix was expressed in HEK cells and immune precipitated with NB611 beads from the cytosolic (C) and membrane (M) fractions. (Left panel) Western blot detecting Alix with mAb 1A12 present in membrane (M) and cytosolic (C) fractions upon immune precipitation with no NB, ᴓ control and NB611. (Right panel) Western blot of the supernatant of the M and C fractions used for IP with NB611 and the corresponding control revealing the presence of Flag-Alix mostly in the cytosolic fraction and its depletion in the membrane fraction upon NB611 IP. Flag-Alix was stained with an anti-Flag antibody.

### NB611 but not NB89 detects Alix recruitment to plasma membrane repair sites

HEK 293 cells knocked-out for Alix (HEK 293 Alix__KO_ cells) (**Figure S6A**) were co-transfected with plasmids encoding either mCherry-Alix and NB611-GFP or mCherry-Alix and NB89-GFP. Recruitment of Alix to the plasma membrane was imaged before and after laser-induced membrane injury. This showed efficient Alix recruitment to injured sites within a few seconds and its colocalization with NB611 (**Figures 3A and C; movies 1 and 2**). In contrast, NB89 displayed only cytosolic staining and does not stain the injured site although Alix was recruited as expected (**Figures 3B and C; movies 3 and 4**). We next transfected HEK293 cells with NB611-GFP and NB89-GFP to demonstrate endogenous Alix recruitment to membrane injury sites. N611-GFP, but not NB89-GFP, stained injury sites (**Figures S6B and C**). Furthermore, NB611-GFP did not localize to the injury sites in HEK 293 Alix__KO_ cells (**Figure S6D**). We therefore conclude that NB611-GFP detects specifically dimeric Alix recruited to injured membranes. Next, we detected CHMP4B recruitment to injury sites in CHMP4B-GFP expressing HeLa cells, starting right after injury and maximal recruitment was observed at 200 sec (**Figure S6E**). Transfection of the CHMP4B-GFP expressing HeLa cells with either NB611-mCherry or NB89-mCherry further showed that NB611-mCherry, but not NB89-mCherry, concentrated at the injured site. Furthermore, both injured sites showed recruitment of CHMP4B-GFP (**Figures S6F and G**) and the kinetics of both CHMP4B and NB611-mCherry recruitment overlapped (**Figures S6H**). We conclude that NB611 detects dimeric Alix at membrane repair sites where CHMP4B is concomitantly recruited.

**Figure 3.**
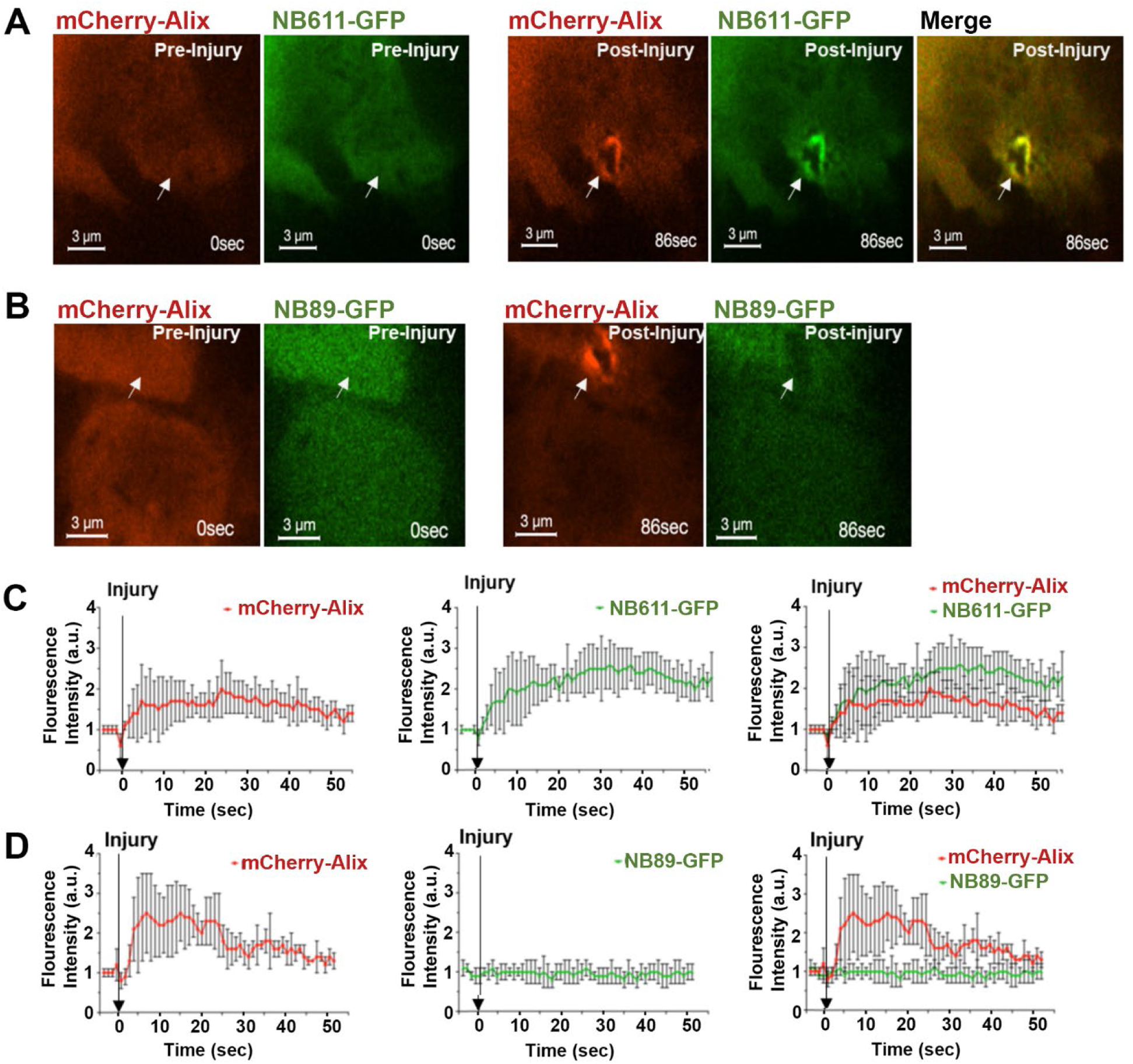
NB611 but not NB89 detects Alix recruitment to plasma membrane repair sites. HEK 293 Alix__KO_ cells were co-transfected with the indicated plasmids and the plasma membrane was injured at 0 sec (indicated by arrows in C and D) using a pulsed laser and imaged by confocal microscopy. **(A)** HEK 293 Alix__KO_ cells were co-transfected with plasmids encoding mCherry-Alix (wild-type) and NB611-GFP. Imaging pre-laser-induced injury (left panels) and post injury (right panels) revealed recruitment of mCherry-Alix together with NB611-GFP and their co-localization at the injured site. White arrows indicate the sites of membrane injury. **(B)** HEK 293 Alix__KO_ cells were co-transfected with mCherry-Alix and NB89-GFP. Images are shown for pre- (left panels) and post-membrane injury (right panels), demonstrating recruitment of mCherry-Alix but not NB89-GFP. This thus indicates that NB89 does not recognize Alix at membrane repair sites. White arrows indicate the sites of membrane injury. **(C)** Graphs showing the fluorescence intensity of mCherry-Alix (left panel) and NB611-GFP (middle panel) recruitment to the sites of membrane injury as shown in A; (right panel) both mCherry-Alix and NB611-GFP recruitment overlap. **(D)** Graphs showing the fluorescence intensity of mCherry-Alix recruitment (left panel), no NB89-GFP recruitment (middle panel) and the overlap of both (right panel) as shown in B at the sites of membrane injury. Each time point corresponds to the fluorescence intensity detected every 5 sec at the site of injury (n=8). Scale bar is 3 µm.

### Mutations that prevent Alix dimerization impair membrane repair

In order to test whether Alix recruitment detected by NB611 allows functional membrane repair, we followed and quantified the TO PRO-3 dye entry after membrane injury. HEK293 cells expressing CHMP4-GFP and mCherry-Alix show accumulation of both CHMP4B and Alix after injury (**Figure 4A, movies 5 and 6**), as expected ^46,47^. This recruitment increased until 325 sec **(Figures 4A and E)**, whereas no measurable increase in TO PRO-3 staining could be observed (**Figure 4A, C and D, movie 7**), suggesting that the membrane has been repaired. In contrast, co-expression of CHMP4B-GFP and of the Alix dimerization mutant mCherry-Alix__mut1_ ^44^ demonstrated CHMP4B and largely diffuse mCherry-Alix__mut1_ recruitment (**Figure 4B; movies 8 and 9**). Appearance of mCherry-Alix__mut1_ at the injury site was largely reduced compared to wild type Alix (**Figures 4 E and F**) and coupled to a steady increase in TO PRO-3 staining (**Figures 4B, C and D; movie 10**), indicative of defective or no membrane repair. We conclude that an active dimeric conformation of Alix is indispensable for membrane repair.

**Figure 4:**
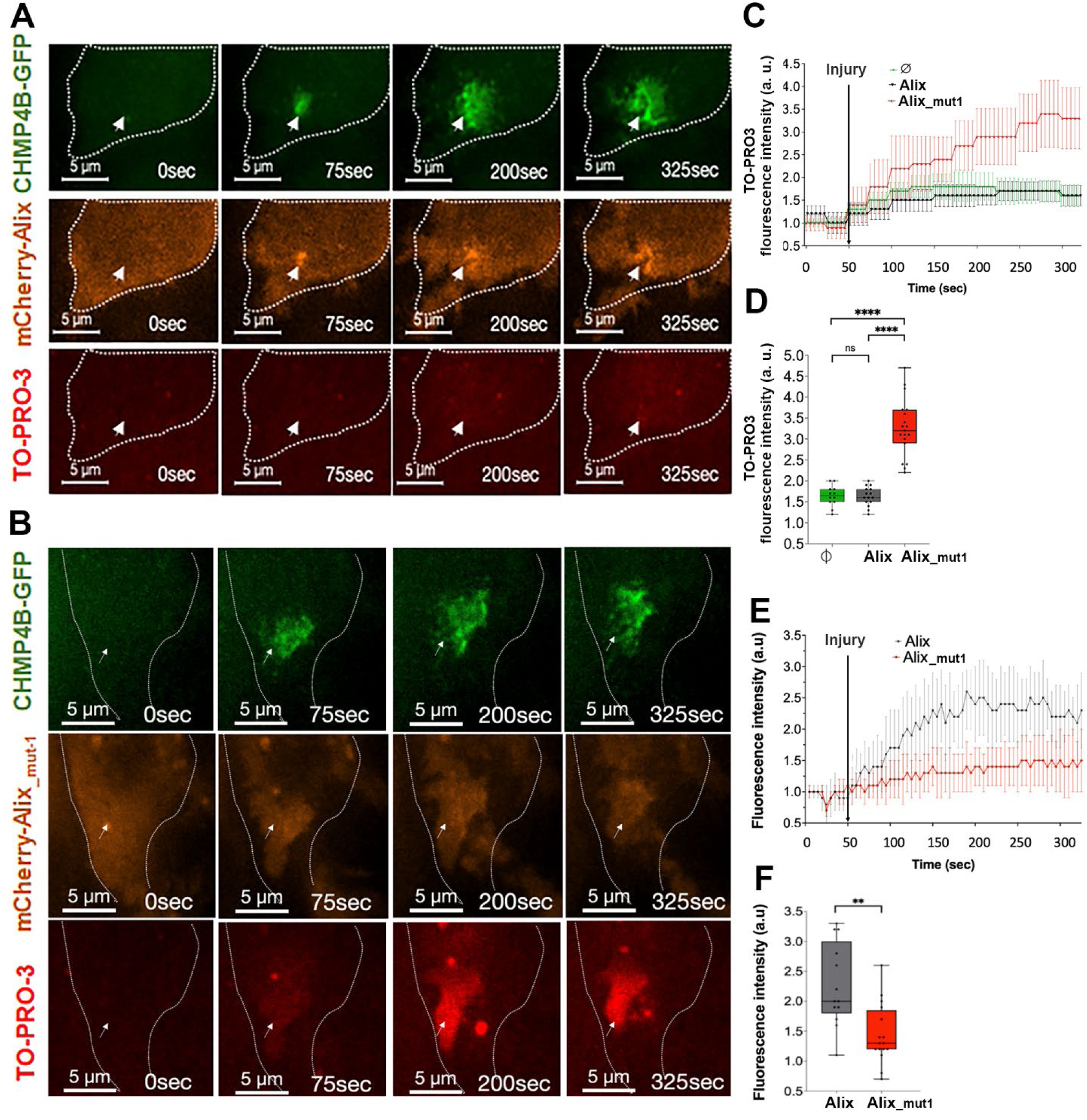
Mutations that prevent Alix dimerization impair membrane repair. HEK293 cells were co-transfected with plasmids encoding the indicated proteins. Plasma membrane was injured using a pulsed laser at time 50 sec and cells were imaged using confocal microscopy. TO PRO-3 dye entry was recorded and quantified after membrane injury. **(A)** Imaging of CHMP4B-GFP (upper panel), mCherry-Alix (middle panel) and TO-PRO-3 dye over 325 sec. **(B)** Imaging of CHMP4B-GFP (upper panel), mCherry-Alix__mut1_ carrying mutations impairing dimerization (middle panel) and TO-PRO-3 dye (bottom panel) over 325 sec. **(C)** The kinetics of TO-PRO-3 dye concentration at the injured plasma membrane is plotted from 0 to 325 sec. The plasma injury is indicated by a black arrow. **(D)** Quantification of the TO-PRO-3 dye intensity at t = 325sec at the injury site, in HEK293 cells transfected with plasmids mCherry-Alix or mCherry-Alix__mut1_. Asterisks indicate statistically significant differences as determined by Welch’s test (n > 10); (****) represents P-value <0.0001. **(E)** Fluorescent intensity of mCherry-Alix and mCherry-Alix__mut1_ at the injured plasma membrane are plotted from 0 to 325 sec. The plasma injury is indicated by a black arrow. **(F)** Quantification of the fluorescence intensity of mCherry-Alix and mCherry-Alix__mut1_ at t = 325 sec at the injury site. Asterisks indicate statistically significant differences as determined by Welch’s test (n > 10); (***) represents P-value <0.001.

### NB611 but not NB89 detects activated Alix at the cytokinetic midbody and abscission site

The cytokinetic midbody at the center of the intercellular bridge (ICB) connecting the dividing daughter cells is a site of active ESCRT-III polymerization ^35,48,49^. Here, Alix plays a key role in the final abscission process, in cooperation with TSG101 ^37,50,51^. Alix appears first as two rings on both sides of the midbody then extends toward the abscission sites, and colocalizes with ESCRT-III polymers at these locations ^35,52–54^. To determine the localization of active Alix during cytokinesis, we stained fixed HeLa cells with either fluorescently labelled NB89 or NB611. Using the same acquisition conditions, NB89 barely labeled the midbody while Alix was present (**Figure 5A and B).** In contrast, NB611 displayed a strong signal at the midbody and toward the abscission site, where it co-localized with Alix (**Figure 5A and B**). NB89 and NB611 signals were drastically diminished upon Alix depletion, showing that they corresponded to Alix (**Figure 5A and B**). Of note, the NB89 signal, although weak, was slightly above background levels. Interestingly, when comparing NB89 (using an enhanced contrast) and NB611 recruitment to the midbody, we found that NB89 recognized a small population of Alix present at the center of the midbody (**Figure 5C**, brackets) whereas NB611 recognized a larger population of Alix localized on the midbody sides (**Figure 5C**, stars). Furthermore, little co-localization of enhanced NB89 was found with CHMP4B at the center of the midbody (**Figure 5D**), whereas extensive co-localization between NB611 and CHMP4B was observed on the sides of the midbody down to the abscission site (**Figure 5E**). This indicates that a minor population of inactive, monomeric Alix is present at the center of the midbody, whereas a major population of dimeric, active Alix revealed by NB611 localizes on the midbody sides and abscission site where it colocalizes with ESCRT-III.

**Figure 5:**
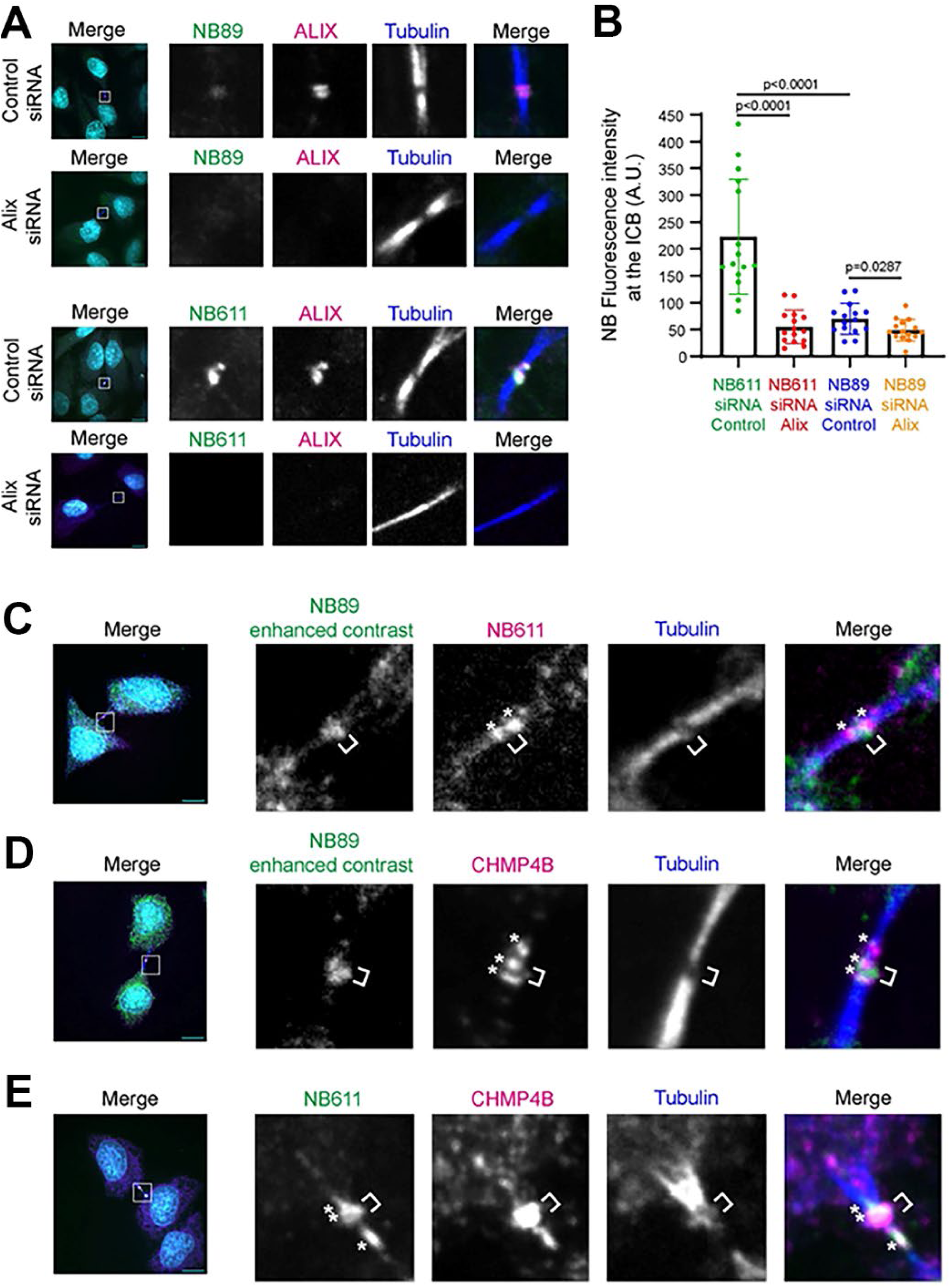
NB611 but not NB89 detects activated Alix at the cytokinetic midbody and abscission site. **(A)** Spinning disk confocal images of HeLa cells treated with indicated siRNAs and labeled with either fluorescently-labelled NB89 or NB611 nanobodies (green), anti-Alix antibody (magenta) and anti-acetylated-tubulin (blue). DAPI in cyan. The acquisition parameters and display for NB89 and NB611 are identical. **(B)** Quantification of the fluorescent intensity (Arb. Unit) of NB89 and NB611nanobodies in HeLa cells treated with either control or Alix siRNAs. Error bars represent SD calculated from 4 independent experiments, each done in triplicate. Two-tailed unpaired Student’s t-test. **(C)** Spinning disk confocal images of HeLa cells labeled with NB89 nanobody (green), NB611 nanobody (magenta) and anti-acetylated-tubulin (blue). DAPI in Cyan. Note that the contrast of the NB89 channel has been enhanced, as compared to (A). **(D)** Spinning disk confocal images of HeLa cells labeled with NB89 nanobody (green), anti CHMP4B antibody (magenta) and anti-acetylated-tubulin (blue). DAPI in Cyan. Note that the contrast of the NB89 channel has been enhanced, as compared to (A). **(E)** Spinning disk confocal images of HeLa cells labeled with NB611 nanobody (green), anti-CHMP4B antibody (magenta) and anti-acetylated-tubulin (blue). DAPI in Cyan. Note that the contrast of the NB89 channel has been enhanced, as compared to (A). Scale bars, 10 μm.

### Dimeric Alix activates CHMP4B polymerization *in vitro*

Numerous *in vitro* studies have shown that CHMP4 and its yeast counterpart Snf7 polymerize spontaneously into filaments depending on their concentration ^44,55–59^. However, under physiological conditions CHMP4B polymerization likely requires a nucleating factor. We used high speed atomic force microscopy (HS-AFM) to test whether Alix in its monomeric or dimeric form can catalyze nucleation of CHMP4B polymerization on supported lipid bilayers (SLB). First, we used HS-AFM to show that at low concentration (500 nM) CHMP4B does not polymerize on SLBs composed of DOPC:DOPS:PIP2 (6:3:1) (**Figure S7A; movie 11**). Similarly, dimeric AlixΔPRD imaging on bilayers with a similar composition did not result in the detection of oligomeric species (**Figure S7B, movie 12**). We next tested combinations of CHMP4B (500 nM) and AlixΔPRD (100 nM). When monomeric AlixΔPRD was used, no CHMP4B polymerization could be detected even after 900 s (**Figure 6A; movie 13**). In contrast, addition of dimeric AlixΔPRD (100 nM) almost instantaneously nucleated CHMP4B polymerization into filaments which could be imaged by HS-AFM, (**Figure 6B**; **movie 14**). Notably, some of the images clearly showed the formation of double filaments most likely coordinated by the two Bro1 domains of dimeric Alix (**Figure 6C and D**). Polymerization of CHMP4B is further favored by the presence of PIP2, as concluded from the observation that the reaction showed reduced oligomer stability on SLBs composed of DOPC:DOPS (6:4) only **(Figure S7C; movie 15**). We conclude that dimeric Alix is a nucleator of CHMP4B polymerization into filaments. Because Alix is recruited to membranes, we propose that the formation of double CHMP4 filaments and their geometry have similar effects on membranes as double CHMP4 filaments coordinated by ESCRT-II, which nucleates CHMP4 via CHMP6 (**Figure 7**), highlighting the importance of double CHMP4 filaments serving as platform for downstream ESCRT-III function.

**Figure 6.**
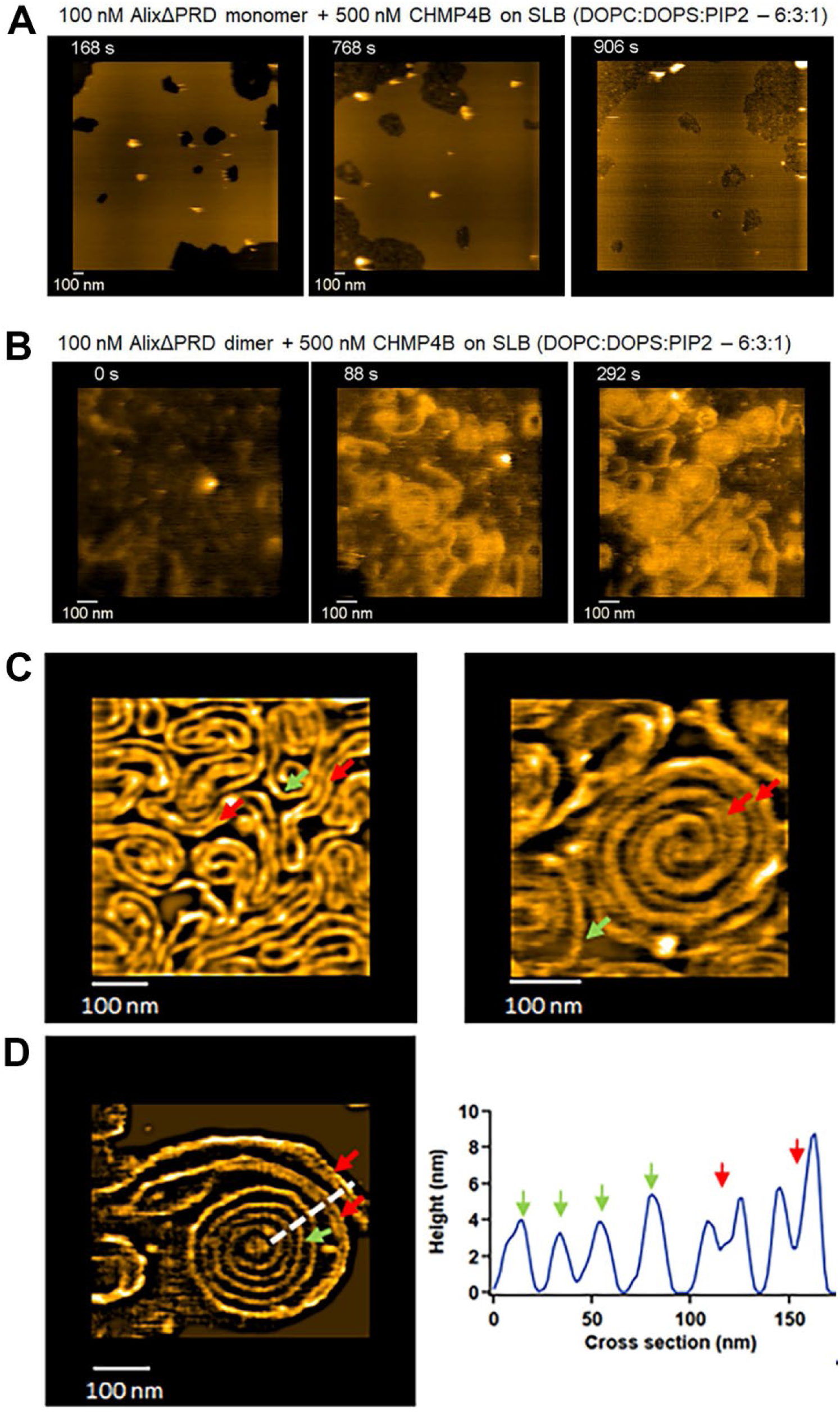
Dimeric Alix nucleates CHMP4B polymerization *in vitro*. **(A)** Time-lapse HS-AFM images of CHMP4B in the presence of monomeric AlixΔPRD show a lack of CHMP4B oligomerisation on supported lipid bilayers; related to **movie 3**. **(B)** Dimeric AlixΔPRD induces the polymerization of stable CHMP4B filaments on supported lipid bilayers containing PIP2; related to **movie 4**. **(C)** Examples of zoomed-in HS-AFM images of dimeric AlixΔPRD induced polymerization (as in **B**) of double filaments on supported lipid bilayers, green and red arrows indicate single and double filaments, respectively. **(D)** Left: example of a CHMP4B spiral with both single (green arrow) and double (Red arrow) filaments assembled in the presence of AlixΔPRD and CHMP4B. Right: cross-section along the white dotted line in the left image, identifying the single (green arrow) and double (red arrow) filaments.

**Figure 7.**
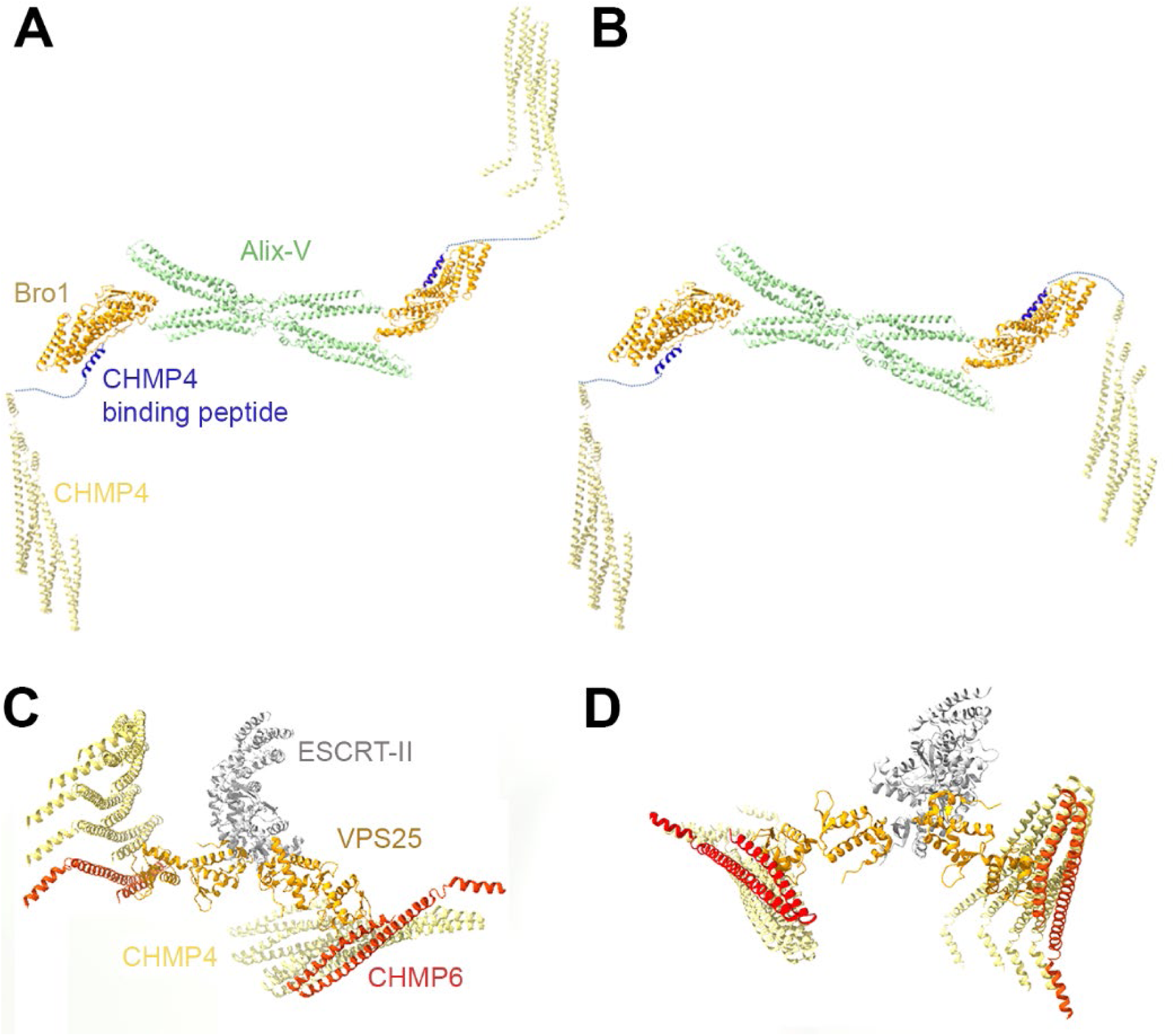
Comparison of models of ESCRT-II in complex with CHMP6-CHMP4 and dimeric Alix in complex with CHMP4. (A and. **B)** Model of dimeric Alix composed of the Bro1 (pdb 2OEV) and the dimeric V-domain (pdb 28LY, this paper). Bro1 interaction with the C-terminal peptide of CHMP4B was modeled based on pdb 3C3Q and the CHMP4B peptide was linked to the open Snf7 (CHMP4) conformation (pdb 5FD7). The orientations of the two CHMP4 filaments coordinated by dimeric Alix could be positioned anti-parallel **(A**) or parallel **(B)**. **(C and D)** Models of the human ESCRT-II complex (pdb 3CUQ) composed of VPS22, VPS36 and two protomers of VPS25. An N-terminal helix of CHMP6 was shown to interact with VPS25 (pdb 3HTU) and its coordinates were used to model CHMP6 in an open conformation in complex with VPS25. CHMP4 protomers (pdb 5FD7) were modeled by binding to CHMP6 mimicking the CHMP4-CHMP4 open conformation interaction. Note that the second winged helix domain of VPS25 interacting with CHMP6 is linked flexibly suggesting that the orientation of the shown CHMP6-CHMP4 polymer could change. Furthermore, based on the model, ESCRT-II can coordinate two filaments polymerized in an antiparallel **(C)** or parallel way **(D).**

## Discussion

Here we demonstrate that the ESCRT-associated scaffolding protein Alix adopts a dimeric conformation associated with ESCRT-catalyzed processes. While Alix dimerization via its V-domain has been shown previously ^30,36,44,45^ our data reveals the mode of dimerization via the exchange of V-domain helix 11, which detaches from arm 1 of the V-shaped structure to interact with arm 1 of the second protomer thereby generating a ‘flat’ X-shaped conformation. It is of note that the individual V-domain arms adopt highly similar conformations as found in monomeric Alix V-domain structures ^30,32,33^. Conformational plasticity is mostly confined to the N-terminal region of arm1 that connects to the Bro1 domain and the hinge region connecting the four V-domain arms. Opening of the monomeric the V-domain via the hinge region that connects two arms has been reported before ^60^ and mutations in the hinge region generated a fully extended open monomeric V domain conformation ^44^. This conformational flexibility is in agreement with the complete opening of the V-domain observed in the dimeric V-domain structure. Molecular dynamics simulations further indicate that the dimeric form remains flexible, albeit less than the monomer. The physiological process that triggers Alix dimerization is yet unknown, but might be regulated by post-translational modifications ^61–63^, ALG-2 interaction ^64–66^ or a combination thereof.

The cytosolic monomeric state of Alix is regulated by its PRD, which keeps Alix in an autoinhibited closed conformation ^41,67^. This is in agreement with results obtained by introducing mutations into the V domain that destabilize its closed conformation leading to increased Alix membrane association and virus budding ^40^. In fact, Alix itself might directly interact with membranes and affect changes in membrane structures ^68,69^. The requirement for Alix activation leading to its dimerization in the ESCRT pathway is further supported by Alix-V and Alix-V-PRD expression that exert a dominant negative effect on HIV-1 budding ^44,45^. This indirectly suggests that truncated Alix participates in dimerization with endogenous Alix and/or in the formation of higher order multimers thereby blocking the ESCRT pathway. Consistent with two conformations of Alix, we detect Alix dimers only in membrane fractions but not in the cytosol of HEK 293 cells, as expected for an ESCRT-catalyzed process.

A major function of Alix in the ESCRT pathway is to recruit ESCRT-III CHMP4. Here we show that dimeric and not monomeric Alix activates CHMP4B to polymerize *in vitro* by employing CHMP4B at a concentration that does not lead to spontaneous CHMP4B (Snf7) filament formation *in vitro* ^44,55,56,58,59^. Our data suggest that dimeric Alix coordinates two CHMP4B filaments that provide the platform for downstream ESCRT-III polymerization of CHMP2A (Vps2) and CHMP3 (Vps24) ^16,18,20^. Likewise, Bro1, the yeast Alix ortholog has been implicated in controlling Snf7 polymerization ^29^. We therefore propose that dimeric Alix functions analogous to ESCRT-II that has two VPS25 (EAP20) protomers ^24^ that recruit CHMP6 ^25^, which in turn engages CHMP4 ^17^ to nucleate its polymerization ^70,71^. Notably, it has been shown in the yeast system that both Vps25 protomers of ESCRT-II are essential for MVB receptor sorting indicating that sorting depends on the geometry of two CHMP4 (Snf7) filaments coordinated by the two Vps25 (VPS25/EAP20) protomers of ESCRT-II ^72^. The comparison of models of dimeric Alix and ESCRT-II shows that both complexes can coordinate two CHMP4 filaments, either in a parallel or anti-parallel way (**Figure 7**). There are, however, two striking differences between ESCRT-II and dimeric Alix regarding the geometry of the coordinated CHMP4 filaments. First, although, the second winged helix domain of ESCRT-II VPS25 is linked flexibly, its direct interaction with the helical hairpin of CHMP6 (Vps20) ^25^ imposes geometrical constraints on the filaments, whereas the C-terminal CHMP4 peptide that interacts with the Alix Bro1 domain is linked flexibly to the core of CHMP4 ^15,31^. Second, the model suggests that ESCRT-II coordinated filaments are spaced by 20 nm versus an ∼35 nm spacing of filaments coordinated by dimeric Alix.

In order to discriminate between the inactive, closed Alix conformation and its activated form *in vivo*, we identified two nanobodies that selectively recognize the two forms. Our data suggest that NB89 binds to the closed state in the cytosol and NB611 to the activated state present in membrane fractions. To further confirm the two states of Alix we used both nanobodies to test Alix recruitment to plasma membrane repair sites since Alix is a major recruiter of ESCRT-III to plasma and lysosomal membrane repair sites ^46,47,73,74^. We found that NB611 but not NB89 detects Alix recruitment to laser-induced plasma membrane injury sites, where in turn CHMP4B is recruited. Ring-like staining of Alix at injury sites is further consistent with Alix multimerization ^36,43,62,75^.

Furthermore, we demonstrate that NB611 interaction with Alix at membrane repair sites does not interfere with membrane repair. Moreover, we found that membrane repair depends on Alix dimerization. Although, the Alix dimerization mutant ^44^ is recruited to the site of membrane repair, the injured membrane is not repaired. This further confirms that Alix dimerization via its V-domain is important to generate the physiological active conformation of Alix in membrane repair. Altogether, these results indicate that NB611 recognizes the active, membrane-associated Alix during plasma membrane repair. Similarly, Alix is known to recruit ESCRT-III at the intercellular bridge during late cytokinetic events leading to abscission ^35,37,48–50,52–54^. The differential staining of NB611 and NB89 show that dimeric Alix (or oligomers of dimeric Alix) is strongly enriched at the midbody sides and at the abscission site, where CHMP4B is actively polymerized ^35,57,76^.

Altogether, this work reveals that dimeric Alix is the conformation of Alix which is active in key cellular ESCRT-III-catalyzed processes. We demonstrate that Alix V-domain-mediated dimerization occurs via domain exchange, which produces an elongated Alix structure that projects two Bro1 domains. Notably, Alix dimers but not monomers nucleate the polymerization of two CHMP4 filaments as shown by HS-AFM. We propose that the double filaments and their geometry is required to coordinate ESCRT-III function during membrane repair and cytokinesis. Importantly, this aligns Alix function with that of ESCRT-II, the other ESCRT-III recruiter that also coordinates two CHMP4 filaments. Although our data reveal that both dimeric Alix and ESCRT-II are central organizers of ESCRT-III assembly, they also explain why ESCRT-II and Alix are not redundant. We propose that these factors generally cannot substitute for one another in ESCRT-mediated membrane remodeling processes because the CHMP4 filaments nucleated by Alix and ESCRT-II differ in spacing and geometry. Nevertheless, the initiation of ESCRT-III polymerization through two CHMP4 filaments that recruit downstream ESCRT-III components is likely a key feature of ESCRT-III function during membrane constriction and fission. This mechanism should therefore be incorporated into future models aimed at understanding the architecture and function of ESCRT-III in membrane remodeling.

## Method Details

### Expression and purification of Alix-V

Alix-V refers to residues 358-714 (https://www.uniprot.org: mouse Q9WU78) having both residues, S568 and S577 mutated to alanine using the QuickChange method (Stratagene). Alix-V was cloned into the bacterial expression vector pETM11.

*Escherichia coli* Rosetta (DE3) cells were transformed with the pETM11-Alix-V plasmid and plated on LB agar (Miller) containing 100 μg/mL kanamycin and chloramphenicol. Expression of the His-tagged Alix-V was induced with 0.1 mM isopropyl-β-D-galactopyranoside (IPTG) at 37°C for 3 h. Cells were harvested by centrifugation at 6,500 rpm for 15 min using a JLA-8.1000 rotor (Beckman Coulter). Pellets were resuspended in 25 mL of buffer A (50 mM Tris-HCl, pH 7.5; 100 mM NaCl; 2 mM β-mercaptoethanol) and lysed by sonication. The lysate was cleared by centrifugation at 20,000 rpm for 20 min using a JA-25.50 rotor (Beckman Coulter), and the supernatant was loaded onto a Ni²⁺-NTA agarose column pre-equilibrated with buffer A. The column was washed with buffer A supplemented with 10 mM imidazole, and proteins were eluted with buffer A containing 250 mM imidazole. The His-tag was removed by overnight cleavage with TEV protease (1:100 w/w, 4°C), followed by dialysis against 4 L of buffer A. To remove the cleaved His-tag, the sample was applied to a second Ni²⁺-affinity column equilibrated in buffer A. The flow-through, containing Alix-V, was collected and dialyzed against 4 L of buffer B (25 mM Bis-Tris, pH 6.3). To separate monomers from dimers Alix-V was applied onto a Mono P column (GE Healthcare), that had been pre-equilibrated with buffer B, and eluted at pH 4.0 using buffer C (100 mL total volume of Pharmalyte: 2 mL of 4-6.5 and 500 μL of 3-10, adjusted to pH 4 with HCl). To prevent precipitation, the eluate was immediately diluted with buffer D (50 mM Tris-HCl, pH 8.8; 100 mM NaCl) until a final pH of 6-7 was achieved. A 10% native polyacrylamide gel was used to assess the oligomeric state and identify fractions containing the dimeric form of Alix-V. These were pooled, concentrated using Amicon Ultra-4 10K MWCO concentrators (Millipore), and further purified by size-exclusion chromatography (SEC) on a Superdex S200 Increase column (GE Healthcare), that had been pre-equilibrated with buffer E (50 mM Tris-HCl, pH 7.5)

### Nanobody discovery, expression and purification

For the discovery of Alix-V specific Nanobodies (NB), two llamas (*Lama glama*) were immunized six times with either a total of 0.9□mg of Alix-V monomers or dimers. The detailed protocols for immunization, nanobody selection, and screening have been described elsewhere ^77^. An immune phage library was generated from mRNA encoding the nanobody open reading frames. Phage display selections were performed on Alix-V monomers immobilized on immunosorbent plates or on Alix-V dimers immobilized on immunosorbent plates in PBS. A screening ELISA was performed on 282 clones using bacterial extracts to identify Alix-V–specific Nanobodies. Colonies yielding a positive ELISA signal were sequenced, and the Nanobody sequences were grouped according to their CDR3 regions into 13 families for the Alix-V monomer and 15 families for the Alix-V dimer. 16 Nanobodies were chosen to be further analyzed and were purified as described elsewhere ^77^.

Nanobody sequences cloned in the vector pMESy4 carrying a PelB signal sequence to direct expression to the periplasmic space ^77^. *Escherichia coli* WK6 (Su-) cells expressing NB611, NB79 or NB 89 were grown in 1L of Terrific Broth supplemented with ampicillin, 0.1% glucose and 2mM MgCl_2_. Protein expression was induced with 1mM IPTG at an OD_600_ = 1-1,2 and cultures were incubated overnight at 28°C. Cultures were harvested by centrifugation (15min at 6000rpm) and pellets were resuspended in 15 ml of buffer F (TES buffer, 0.2 M Tris pH 8; 0.5 mM EDTA; 0.5 M sucrose) per liter of culture and gently stirred for at least 1h at 4 °C. Subsequently, 30 ml of 4-fold diluted TES buffer per 1L culture of cell pellet was added and the suspension was incubated with gentle shaking for at least 45 min at 4 °C. The suspensions were centrifuged for 30 min at 8,000 rpm, and the supernatant corresponding to the periplasmic fraction was collected. The periplasmic extract was loaded onto Ni-NTA resin pre-equilibrated with buffer G (50 mM Na phosphate, 1M NaCl, pH 7.0) and washed with buffer H (50 mM Na phosphate, 1M NaCl, pH 6.0). Proteins were eluted with buffer G containing 250 mM imidazole. Final purification was performed by size-exclusion chromatography using a Superdex 200 Increase column (GE Healthcare) equilibrated in HBS buffer I (25 mM HEPES pH7.5, 150mM NaCl).

For mammalian expression nanobody sequences of NB89 and NB611 were cloned into expression vectors pEGFP-N1 (Clonetech) and pEmCherry-N1 (vector pEGFP-N1 with GFP replaced by mCherry) to generate expression plasmids NB89-mCherry, NB89-GFP, NB611-mCherry and NB611-GFP employed in the membrane repair assays.

### Expression and purification of full-length Alix

HEK293FS cells were cultured in Freestyle 293 medium (Gibco) and maintained at 37°C in a 10% CO₂ atmosphere, under agitation at 150 rpm. On the day of transfection, a 300 mL suspension culture at a density of 1 × 10□ cells/mL was freshly prepared. For transfection, 300 µg of pCi Flag-mAlix WT plasmid DNA were diluted in 15 mL of Opti-MEM (Gibco), and 900 µL of 1 mg/mL linear 25 kDa PEI (Polysciences) were added dropwise. The DNA-PEI mixture was incubated for 20 mins at room temperature and then added directly to the cell culture. After 72 hours of expression, cells were harvested by centrifugation at 264 × g for 10 mins, washed once with 1× PBS, and lysed in 75 mL of lysis buffer J (20 mM Tris-HCl pH 7.4,150 mM NaCl, 1 mM MgCl₂, 1% Triton X-100; EDTA-free protease inhibitors (Roche)). The lysate was incubated on ice for 15 mins, then clarified by centrifugation at 16,000 × g for 10 mins using a JA-25.50 rotor (Beckman Coulter). The supernatant was loaded onto an anti-FLAG affinity column (sigma) pre-equilibrated with buffer A (50 mM Tris-HCl pH 7.5; 100 mM NaCl; 2 mM β-mercaptoethanol). Bound proteins were eluted using 100 mM glycine, pH 3,5. To immediately neutralize the eluate, fractions were collected into tubes pre-filled with buffer D (50 mM Tris-HCl pH 8.8; 100 mM NaCl). As a final purification step, the sample was subjected to size-exclusion chromatography using a Superdex S200 Increase column (GE Healthcare) pre-equilibrated with HBS buffer I (25 mM HEPES pH 7.5 ;150 mM NaCl)

### Bio Layer Interferometry

Nanobodies were biotinylated following the protocol provided in the ForteBIO Protein Biotinylation Kit for Immobilization on Streptavidin Biosensors. Excess, unbound biotinwas removed by desalting the nanobodies using Zeba Spin Desalting Columns (Thermo Scientific). Streptavidin biosensors were coated with biotinylated nanobodies at final concentration of 50 µg/mL, prepared by diluting the protein in HBS buffer (25mM HEPES pH 7.5, 150mM NaCl, supplemented with 0.02% Tween-20). A threshold capture of 1.2 nm was applied. Alix concentration series were prepared in buffer A according to the construct and nanobody used, as follows:

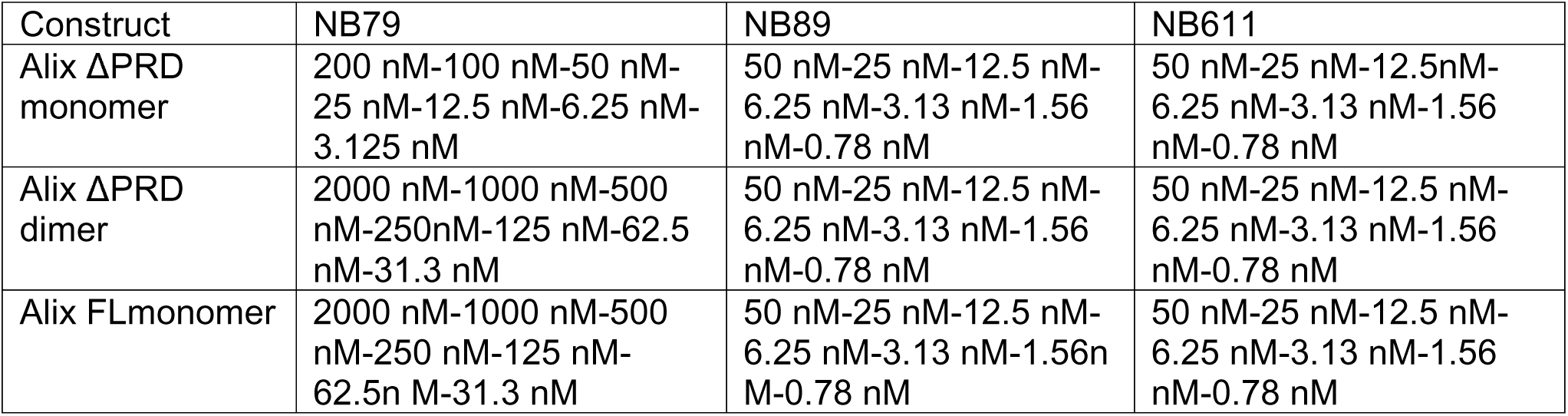

| Construct | NB79 | NB89 | NB611 |
| --- | --- | --- | --- |
| Alix ΔPRD monomer | 200 nM-100 nM-50 nM-25 nM-12.5 nM-6.25 nM-3.125 nM | 50 nM-25 nM-12.5 nM-6.25 nM-3.13 nM-1.56 nM-0.78 nM | 50 nM-25 nM-12.5 nM-6.25 nM-3.13 nM-1.56 nM-0.78 nM |
| Alix ΔPRD dimer | 2000 nM-1000 nM-500 nM-250 nM-125 nM-62.5 nM-31.3 nM | 50 nM-25 nM-12.5 nM-6.25 nM-3.13 nM-1.56 nM-0.78 nM | 50 nM-25 nM-12.5 nM-6.25 nM-3.13 nM-1.56 nM-0.78 nM |
| Alix FLmonomer | 2000 nM-1000 nM-500 nM-250 nM-125 nM-62.5 nM-31.3 nM | 50 nM-25 nM-12.5 nM-6.25 nM-3.13 nM-1.56 nM-0.78 nM | 50 nM-25 nM-12.5 nM-6.25 nM-3.13 nM-1.56 nM-0.78 nM |

All measurements were performed in triplicate. Data were analyzed using the ForteBio software. For the AlixΔPRD constructs (both monomeric and dimeric forms), a 1:1 binding model was applied. For the full-length Alix construct, a heterogenous ligand binding model was used.

### Alix-V-NB79 complex formation and crystallization

Alix-V and NB79 were mixed in a 1:2 molar ratio for 1h at room temperature prior to purification by SEC using a Superdex 200 10/300GL increase column in buffer E (50 mM Tris pH 7.5). Fractions from the major peak were shown to contain both Alix-V and NB79. Fractions were concentrated to 12 mg/ml and used for crystallization screening on the ISBG HTX crystallization platform. Initial crystallization conditions were refined manually and diffracting crystals were obtained by micro seeding in 0.1 M HEPES, pH 7.5, 8% (w/v) PEG 8000, 8% (v/v) ethylene glycol and 0.01 M sodium acetate, pH 4.6. Crystals were grown using the hanging drop vapor diffusion method at 20°C. Single crystals were mounted in cryo-loops and flash cooled in liquid nitrogen. X-ray diffraction data were collected under a nitrogen stream at 100 K at the European Synchrotron Facility (ESRF), Grenoble, France.

### Structure determination and refinement

Diffraction data were collected to 2.67 Å on beamline MASSIF-1 ^78^ (ESRF). The diffracting crystal was in space group P2_1_ and displayed two 1:1 Alix-V-NB79 complexes per asymmetric unit. Statistics on data collection and refinement are summarized in Table S1. X-ray diffraction images were indexed and scaled with the XDS program package ^79^. ADXV (https://www.scripps.edu/tainer/arvai/adxv.html) and XDSGUI ^80^ were used to check data quality and resolution cutoff ^81–85^. The anisotropic diffraction limit was determined using the STARANISO server (http://staraniso.globalphasing.org/cgi-bin/staraniso.cgi). The reduced X-ray diffraction data was imported into the CCP4 program suite ^86^. The Alix-V-NB79 structure was solved by molecular replacement using PHASER ^87^. The Alix (2OJQ) and nanobody (6QUP) model chains were placed sequentially. Unit cell macromolecular assembly was determine using the PISA server ^88^. The structure was completed by cycles of manual model building with COOT ^89^. Water molecules were added to the residual electron density map as implemented in COOT ^90^. Crystallographic macromolecular refinement was performed with REFMAC ^91^. Cycles of model building and refinement were performed until *R_work_* and *R_free_* converged ^92,93^. The TLS definition ^94^ was determined and validated using the TLSMD ^95^ and PARVATI ^96^ servers. The stereochemical quality of the refined models was verified with MolProbity ^97^, PROCHECK ^98^ and PDB-REDO ^99^. Secondary structure assignment was performed by DSSP ^100^ and STRIDE ^101^.

### SAXS data collection and analyses

Small-angle X-ray scattering (SAXS) of wild type Alix were collected at the BioSAXS beamline (SWING) of the SOLEIL synchrotron. 12 mg/ml of protein were separated on a Superdex Increase 200 5/150 GL column (Cytiva) in buffer E (50mM Tris HCl pH7.5). SAXS measurements were performed every second with an Eiger 4□M detector (distance of 2.00□m) allowing a q range of 0.004 to 0.55□Å-1 with a wavelength of 1.03□Å. Data analyses were performed using PRIMUS from the ATSAS package ^102^. For data reduction, buffer acquisition was performed before and after each sample measurement. Distances P(r), Dmax and radius of gyration (*R* _G_) were determined from the Guinier approximation. All preparations analyzed were monodisperse, as evidenced by linearity in the Guinier region of the scattering data and agreement of the I_0_ and R_g_ values determined with inverse Fourier transform analyses using GNOM ^103^. The *ab initio* models (low-resolution envelope of the protein) were calculated with DAMMIN ^104^. 20 independent envelopes were averaged by DAMAVER ^105^. Finally, the final envelope was fit against the scattering data using DAMMIF ^106^.

### HS-AFM imaging and analysis

AFM images were acquired in amplitude modulation tapping mode in liquid, using high-speed atomic force microscopes (RIBM, Japan) ^107,108^. The HS-AFM imaging was performed using USC-F1.2-k0.15 cantilevers (NanoWorld, Switzerland), an ultrashort cantilever with a nominal spring constant of 0.15 N/m and a resonance frequency ≈ 0.5 MHz in liquid. All HS-AFM recordings were performed at room temperature in a buffer chamber containing a maximum of 100 µl of HBS buffer (25 mM HEPES pH 7.5 ;150 mM NaCl). Supported lipid bilayers (SLB) were created by fusing large unilamellar vesicles (LUVs) containing DOPC: DOPS: PIP2 at a 6:3:1 molar ratio on mica (freshly cleaved) for 15 mins. SLBs without PIP2 were formed of DOPC: DOPS at a 60:40 molar ratio. After incubation, the surface was gently washed to remove free and unfused vesicles. HS-AFM imaging was started in order to localize the SLB patches. Once a patch was found, CHMP4B and/or Alix were sequentially added to the required concentration, as mentioned in the results section, while maintaining continuous imaging. All reported HS-AFM images were analyzed using Igor Pro (RIBM script) and ImageJ with additional home-written plugins ^109^ with tilt and contrast corrections. All experiments reported in the manuscript were repeated at least three times.

### Cells and cell culture

HEK293 cells (ATCC CRL-1573), HeLa (ATCC CCL-2) cell line and HeLa cells expressing CHMP4B-LAP-eGFP (CHMP4B-GFP) ^35^ were grown in Dulbecco’s Modified Eagle Medium (DMEM) GlutaMax (31966; Gibco, Invitrogen Life Technologies) supplemented with 10 % fetal bovine serum in 5% CO_2_ at 37□°C.

### Membrane repair assay

For laser-induced membrane injury experiments, HEK293 cells were seeded on 8-well glass LabTek chambers pre-coated with 10µg/ml of polyornithine, in DMEM containing 10% FBS. Cells were transfected with 50 ng per well with plasmids NB611-mCherry, NB611-GFP, NB89-mCherry, NB89-GFP or 15 ng per well with CHMP4B-LAP-eGFP plasmids. 20h post transfection, the medium was replaced with phenol red-free DMEM containing 10% FBS.

Live cell imaging was performed using a spinning disk confocal Olympus IX81/Yokogawa CSU-X1 microscope, equipped with an Andor iXon EMCCD camera, a GATACA iPulse laser bench, and iLAS2 FRAP device. The setup is equipped with a CO_2_/temperature-controlled chamber set at 37°C; 5% CO_2_. Images were acquired every 5 seconds under continuous focus control (ZDC: Zero Drift Control device) with a 100x oil immersion apochromatic objective (Olympus UAPON100X, NA1.49). After the first 5 frames, a manually defined area of approximately 530 x 660 nm was ablated for 40 ms using a 355 nm pulsed laser at 20% of its maximum power (Teem Photonics, 400ps pulses at 20kHz - 2kW) for 20 repetitions. Recovery is monitored over 100 frames (∼8.5 mins). Before each experiment, a test ablation was performed to ensure correct targeting, power and optimal focus. Microscope control was performed with Metamorph (Molecular Device™) and image processing was performed using Imaris. To assess plasma membrane integrity, cells were incubated with TO-PRO-3 iodide (ThermoFisher; 0.3□µM final concentration) in imaging buffer. TO-PRO-3 is a cell-impermeant DNA dye that fluorescens in the far-red channel (excitation: 642 nm; emission: 661 nm) only upon membrane compromise. *Statistics.* Plots and statistical test (Welch’s test) were performed using GraphPad Prism software version 11. The presented values are displayed as mean ± SD n > 10 cells per conditions. *P*-values are indicated in the figures. Welch’s test (n > 10); (****) represents P-value <0.0001.

### Immunoprecipitation from cytosolic and membrane fractions

HEK293 cells were transfected with plasmids Flag Alix FL pcDNA. Cells were washed with PBS before being harvested. The homogenate was centrifuged at 500 x g for 5min and the pellet was resuspended in buffer A (50 mM Tris HCl pH 7.5, 1 mM MgCl_2_, 1 mM PMSF and complete EDTA-protease inhibitors (Roche)). The suspension was sonicated for 10 s to disrupt the cells. The sample was then centrifugated at 1,500 × g for 5 min at 4°C to pellet nuclei and debris and the supernatant was subsequently centrifuged at 20.000 × g for 1 h at 4°C. The resulting supernatant was collected as the cytosolic fraction, and the pellet (membrane fraction) was resuspended in membrane lysis buffer A supplemented with 0.5% Triton X-100 and incubated on ice for 15min. Cytosolic and membrane fractions were incubated with 15 µg of purified nanobody NB611 for 1 h at 4°C on a rotating platform. Samples were then incubated with protein A-conjugated beads (pre-blocked with 5% BSA; 0.5% triton) for 1 h at 4°C on a rotating platform. Beads were washed three times with lysis buffer to remove unbound protein. Bound proteins were eluted by incubating beads with 2× Laemmli buffer for 5 mins at 95°C. Eluted proteins were separated by SDS-PAGE and transferred onto a PVDF membrane (Millipore). Membranes were blocked with 5% non-fat dry milk in PBS-Tween for 30 min at room temperature and incubated 1 h with the anti-Alix antibody 1A12 (1:1000)(Santa Cruz). The membrane was then incubated with anti-mouse IgG conjugated to HRP (1:10 000, Jackson ImmunoResearch) in PBS-Tween-milk solution for 1h at room temperature. Detection was performed using enhanced chemiluminescence (ECL) substrate (Thermo Scientific) and visualized using a ChemiDoc imaging system (Bio-Rad).

### Co-Immunoprecipitation of Flag-Alix and Myc-Alix

HEK293 cells were transfected with pcDNA-Flag-Alix (full length Alix) and pcDNA-Myc-Alix plasmids. Cells were washed with PBS before being harvested by scraping. The homogenate was centrifuged at 500 x g for 5 min and the pellet was resuspended in buffer K (50 mM Tris HCl pH7.5, 1 mM MgCl_2_, 1 mM PMSF and Complete EDTA-protease inhibitors (Roche)). The suspension was sonicated for 10 sec to disrupt the cells. After sonication, samples were centrifugated at 1,500 × g for 5 min at 4°C to pellet nuclei and debris. The supernatant was then centrifuged at 20,000 × g for 1 h at 4°C. The resulting supernatant was collected as the cytosolic fraction, and the pellet (membrane fraction) was resuspended in membrane lysis buffer K supplemented with 0.5% Triton X-100 and incubated on ice for 15min.

Cytosolic and membrane fractions were incubated with anti-flag M2 affinity gel (sigma Aldrich-A2220) or Anti-c-Myc agarose (Pierce-20168) pre-blocked with 5% BSA, 0.5% Triton for 1 h at 4°C with gentle rotation. Beads were washed three times with lysis buffer K to remove unbound material. Bound proteins were eluted by incubating beads with 2× Laemmli buffer for 5 mins at 95°C. Eluted proteins were separated by SDS-PAGE and transferred onto a PVDF membrane (Millipore). Membranes were blocked with 5% non-fat dry milk in PBS-Tween for 30min at room temperature and incubated 1hr with rabbit anti-myc (Sigma Aldrich) or rabbit anti-flag (Sigma Aldrich F7425) antibody. Membranes were further incubated for 1 h with secondary antibodies coupled to HRP. Detection was performed using enhanced chemiluminescence (ECL) substrate (Thermo Scientific) and visualized using a ChemiDoc imaging system (Bio-Rad).

### NB611 and NB89 midbody staining

For silencing experiments, HeLa cells were transfected with 25 nM siRNAs (siControl or siAlix) for 5 days using Lipofectamine RNAiMAX (Invitrogen), following the manufacturer’s instructions. Control non-targeting siRNA (5’UGGUUUACAUGUUGUGUGA[dU][dU]3′), Alix siRNA (5’CCUGGAUAAU GAUGAAGGA[dU][dU]3′).

Immunofluorescence and image acquisition was performed by spinning disk confocal microscopy. Cells were grown on coverslips and fixed with Methanol for 5□min at −20°C. Cells were then permeabilized and blocked with PBS containing 0.2% bovine serum albumin (BSA) and 0.1% triton for 10 min and blocked with PBS containing 0.2% BSA for 30 min. Cells were then successively incubated for 1□h at room temperature with primary human anti-acetylated-tubulin (C3B9-hFc, A-R-H#39) and either with mouse anti-Alix (Biolegend, 634502) or rabbit anti-CHMP4B (Proteintech, 13683-1-AP). Then, cells were incubated for 2 hours and 30 min with nanobody anti-Alix (NB89 and NB611, stained with alexa-488 or alexa-555, 1:250 dilution) and secondary antibodies (1:1000, Jackson Laboratory) diluted in PBS containing 0.2% BSA, before DAPI staining for 5min (0.5□mg/mL, Serva). Cells were mounted in Fluoromount G (Southern Biotech). Images were acquired with an inverted Nikon Eclipse Ti-E microscope equipped with a CSU-X1 spinning disk confocal scanning unit (Yokogawa) coupled to a Prime 95S scientific complementary metal-oxide semiconductor (sCMOS) camera (Teledyne Photometrics). Plots and statistical test (unpaired student t test) were performed using GraphPad Prism. The presented values are displayed as mean ± SD from 15 cells per conditions. *P*-values are indicated in the figures. *P*-value > 0.05 was considered as nonsignificant.

### Theory and computations

#### Computational assays

The first assay consisted of dimeric Alix-V (pdb 28LY, this study) immersed in an aqueous medium enclosed in a rhombohedral cell of initial dimensions equal to 70 × 110 × 230 Å^3^. To prevent tumbling of the protein dimer, the principal axis of inertia of the three-helix bundle formed by residues 430-470, 480-523 and 643-708 of the first monomer was aligned with the long edge of the simulation cell, thereby allowing a lesser amount of water molecules to be used. The second assay consisted of AlixΔPRD (pdb 2OEV) immersed in an aqueous medium enclosed in a cubic cell of initial dimensions equal to 170 × 170 × 170 Å^3^. All computational assays were electrically neutral, with a sodium chloride concentration set to 150 mM.

#### Equilibrium simulations

All MD simulations were carried out with NAMD 3.0 ^110^. The CHARMM36m force field was employed to represent proteins and ions ^111,112^, whereas water molecules followed the TIP3P description ^113^. The MD trajectories were generated in the isothermal–isobaric ensemble at 300 K and 1.013 bar. Temperature control relied on the stochastic velocity rescaling algorithm ^114^, and pressure regulation was achieved with the Langevin piston approach ^115^. Periodic boundary conditions were applied. Short-range electrostatic and van der Waals interactions were truncated smoothly with a 12-Å cutoff. Long-range electrostatic interactions were evaluated through the particle–mesh Ewald method, with a grid spacing of 1.2 Å ^116^. Hydrogen mass repartitioning ^117^ was applied in every simulation, enabling the use of a 4 fs time step. The integration scheme followed the r-RESPA multiple time-step algorithm ^118^, with long-range forces updated every 8 fs, and short-range forces every 4 fs. All covalent bonds involving hydrogen atoms were constrained using SHAKE/RATTLE ^119,120^, while SETTLE ^121^ was employed to maintain water molecules in their equilibrium geometry.

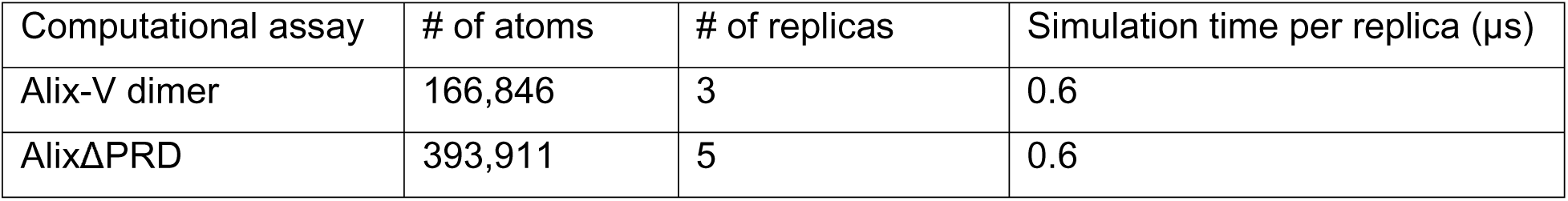

| Computational assay | # of atoms | # of replicas | Simulation time per replica (μs) |
| --- | --- | --- | --- |
| Alix-V dimer | 166,846 | 3 | 0.6 |
| AlixΔPRD | 393,911 | 5 | 0.6 |

Solvated dimeric Alix-V and AlixΔPRD were simulated in triplicate and quintuplicate, which corresponds to an aggregate time of 1.8 and 3.0 µs, respectively. Each simulation was prefaced by an energy minimization, a 10-ns thermalization with strong positional harmonic restraints (force constant of 10 kcal/mol·Å^2^) on the protein heavy atoms, a 10-ns thermalization with soft positional harmonic restraints (force constant of 1 kcal/mol·Å^2^) on the protein backbone atoms, and a 20-ns equilibration devoid of geometric restraints. The analyses of the MD trajectories were performed using the collective variables library for molecular simulation and analysis programs Colvars ^122^ and the visualization program VMD ^123^.

### Figure Generation

Molecular graphics figures were generated with PyMOL (W. Delano; The PyMOL Molecular Graphics System, Version 1.8 Schrödinger, LLC, (http://www.pymol.org). The models shown in Figure 7 were assembled with UCSF ChimeraX ^124^. Sequence alignments were performed with Clustal Omega ^125^ and ESPript ^126^.

## Data Availability

Coordinates and structure factors were deposited in the Protein Data Bank (https://www.ebi.ac.uk/pdbe/), pdb accession code 28LY. Sequences of nanobodies were deposited in the NanoSaurus database (https://www.nanosaurus.org/), accession codes SD-GTNB (NB89), SD-WM6I (NB611) and SD-DYP1 (NB79).

## Acknowledgement

WW acknowledges support from the ‘Institut Universitaire de France’ (IUF), the ‘Fondation pour la Recherche Medicale (FRM, Equipe FRM 2023 202303016333) and the ‘Agence Nationale pour la Recherche’ (ANR-23-CE11-0008-01). AE acknowledges support from Institut Pasteur, CNRS, the ‘Fondation pour la Recherche Medicale’ (FRM, Equipe FRM 202103012627 and 202503020073) and the ‘Agence Nationale pour la Recherche’ (ANR-23-CE11-0008-02). EP and JS acknowledge the support and the use of resources of Instruct-ERIC (PID9044), part of the European Strategy Forum on Research Infrastructures (ESFRI), and the Research Foundation - Flanders (FWO) for their support to the Nanobody discovery (VID16340). WW acknowledges access to the platforms of the Grenoble Instruct-ERIC center (IBS and ISBG; UAR 3518 CNRS-CEA-UGA-EMBL) within the Grenoble Partnership for Structural Biology (PSB), with support from FRISBI (ANR-10-INBS-05-02) and GRAL, a project of the University Grenoble Alpes graduate school (Ecoles Universitaires de Recherche) CBH-EUR-GS (ANR-17-EURE-0003). The IBS acknowledges integration into the Interdisciplinary Research Institute of Grenoble (IRIG, CEA) and financial support from CEA, CNRS and UGA. We thank the ESRF-EMBL Joint Structural Biology Group for access and support at the ESRF beam lines, Jose Marquez (EMBL) from the HTX crystallization facility, Caroline Mas and Jean-Baptiste Reiser for assistance on IBS-ISBG platforms and Katleen Willibal for technical assistance during Nanobody discovery.

## Author contributions

W.W. designed and coordinated the research, interpreted experiments, supervised and received funding for the study. W.W., R.S; W.H.R., A.D., A.E. and J.S. secured funding and supervised personnel. N.M., C.C.-M., S.M., E.P., S.F., P.M., C.Cha., J.-M.B., J.-P.K. and C.Chi. performed research and analyzed data. W.W. wrote the paper with input from all authors.

