## Supplemental Figures for "Dimeric Alix nucleates ESCRT-III CHMP4 polymerization"

### Supplemental tables and figures

| Ligand | Analyte | $K_D$ (M) | $k_a$ (1/Ms) | $k_d$ (1/s) |
|---|---|---|---|---|
| **NB79** | Alix$\Delta$PRD$_{Monomer}$ | $1,5.10^{-8}$ ± 0,32 | $4.10^{-4}$ ± 1,35 | $5,3.10^{-4}$ ± 1,24 |
| | Alix$\Delta$PRD$_{Dimer}$ | $1,2.10^{-7}$ ± 0,17 | $8,5.10^{-3}$ ± 2,35 | $0,9.10^{-3}$ ± 0,15 |
| **NB89** | Alix$\Delta$PRD$_{Monomer}$ | $1,6.10^{-9}$ ± 0,56 | $3,3.10^{-5}$ ± 0,67 | $4,4.10^{-4}$ ± 0.76 |
| | Alix$\Delta$PRD$_{Dimer}$ | $2,4.10^{-9}$ ± 0.53 | $3,1.10^{-5}$ ± 0.98 | $6,9.10^{-4}$ ± 1.93 |
| **NB611** | Alix$\Delta$PRD$_{Monomer}$ | $3,5.10^{-9}$ ± 1.51 | $4,8.10^{-5}$ ± 2.67 | $4,2.10^{-3}$ ± 3,11 |
| | Alix$\Delta$PRD$_{Dimer}$ | $1,7.10^{-9}$ ± 0.71 | $1,8.10^{-5}$ ± 0.14 | $3,2.10^{-4}$ ± 0.92 |

**Table S1: Biolayer interferometry (BLI analyses.** Rate constants for association ($k_a$) and dissociation ($k_d$) and equilibrium dissociation constants ($K_D$) were determined for evaluating the binding affinity of NB79, NB89 and NB611 to the different Alix constructs as indicated. N=3; mean ± SD.

**Table S2: Crystallographic data collection, phasing and refinement statistics.**

| | |
|---|---|
| **Data collection** | ESRF ID30A-1, Pilatus 2M detector) |
| Space group | $P2_1$ |
| Wavelength | 0.96546 |
| Unit cell a, b, c (Å) | 84.26, 70.87, 130.61 |
| α, β, γ (°) | 90°, 108.81°, 90° |
| Overall resolution (Å) | 45.3-2.67 |
| High resolution shell (Å) | 2.82-2.66 |
| Nb observed/unique reflexions | 66927/21902 |
| Multiplicity | 3.05 |
| Completeness (%) | 90.5 (66.7) |
| Mosaicity (°) | 0.162 |
| $R_{sym}$ (last shell) | 6.7 (75.3) |
| $R_{pim}$ | 4.5 (72.9) |
| I/σ(I) (last shell) | 8.7 (1.7) |
| Wilson plot B-factor (Å$^2$) | 118.416 |
| CC (1/2) | 99.5 (47.3) |
| **Molecular replacement** | |
| Phaser LLG | 1006.96 |
| $R_{work}$/$R_{free}$ (%) | 37.28/44.87 |
| **Refinement** | |
| Resolution (Å) | 45.30-2.67 |
| No. reflections | 19891 |
| Reflections used for $R_{free}$ | 2011 |
| $R_{work}$ / $R_{free}$ | 20.30/25.90 |
| No. atoms | 7114 |
| **Stereochemistry** | |
| RMS deviation, bond lengths (Å) | 0.006 |
| RMS deviation, bond angles (º) | 1.288 |
| Mean B-factor (Å$^2$) | 85.353 |
| Residues in most favored/allowed region of Ramachandran plot (%) | 100 |
| PDB code | 28LY |

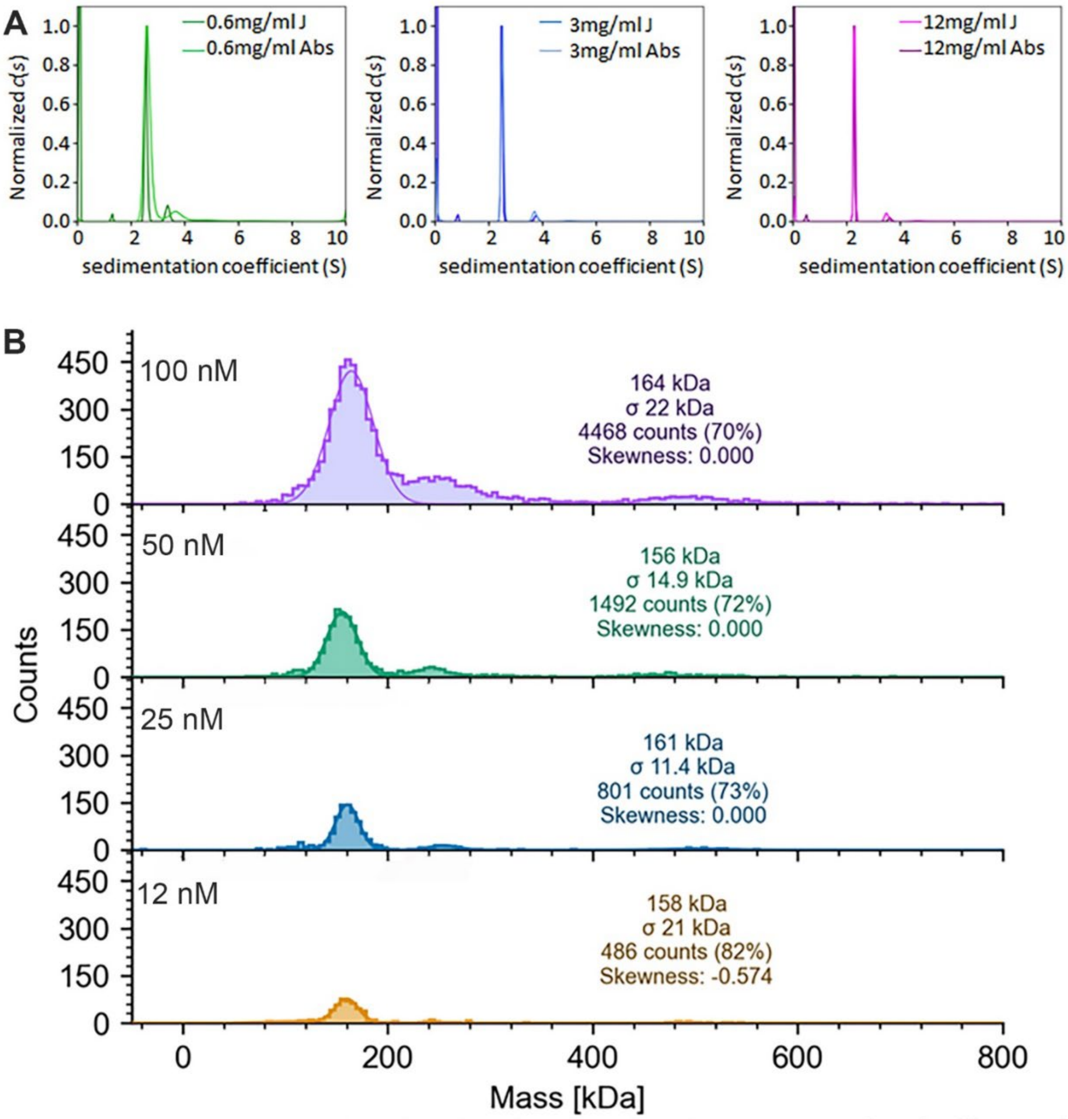


**Figure S1. Characterization of monomeric and dimeric Alix**

**(A)** Monomeric AlixΔPRD was analyzed by analytical ultracentrifugation at 0.6 mg/ml (7.5 μM), 3.0 mg/ml (37.0 μM), and 12 mg/ml (150,0 μM). The main peak at a sedimentation coefficient (S) of 2,2 accounts for 86 ± 4% of the total signal and corresponds to AlixΔPRD monomers (f/fmin = 1.08, theoretical Mw = 79.9 kDa) indicating no concentration-dependent dimerization. The peaks detected by interference (J) and absorbance (Abs) overlap.

**(B)** Dimeric AlixΔPRD was analyzed by mass photometry at concentrations of 100, 50, 25 and 12 nM, which revealed consistent molecular masses of ~160 kDa in agreement with its theoretical dimeric molecular weight of 159.9 kDa.

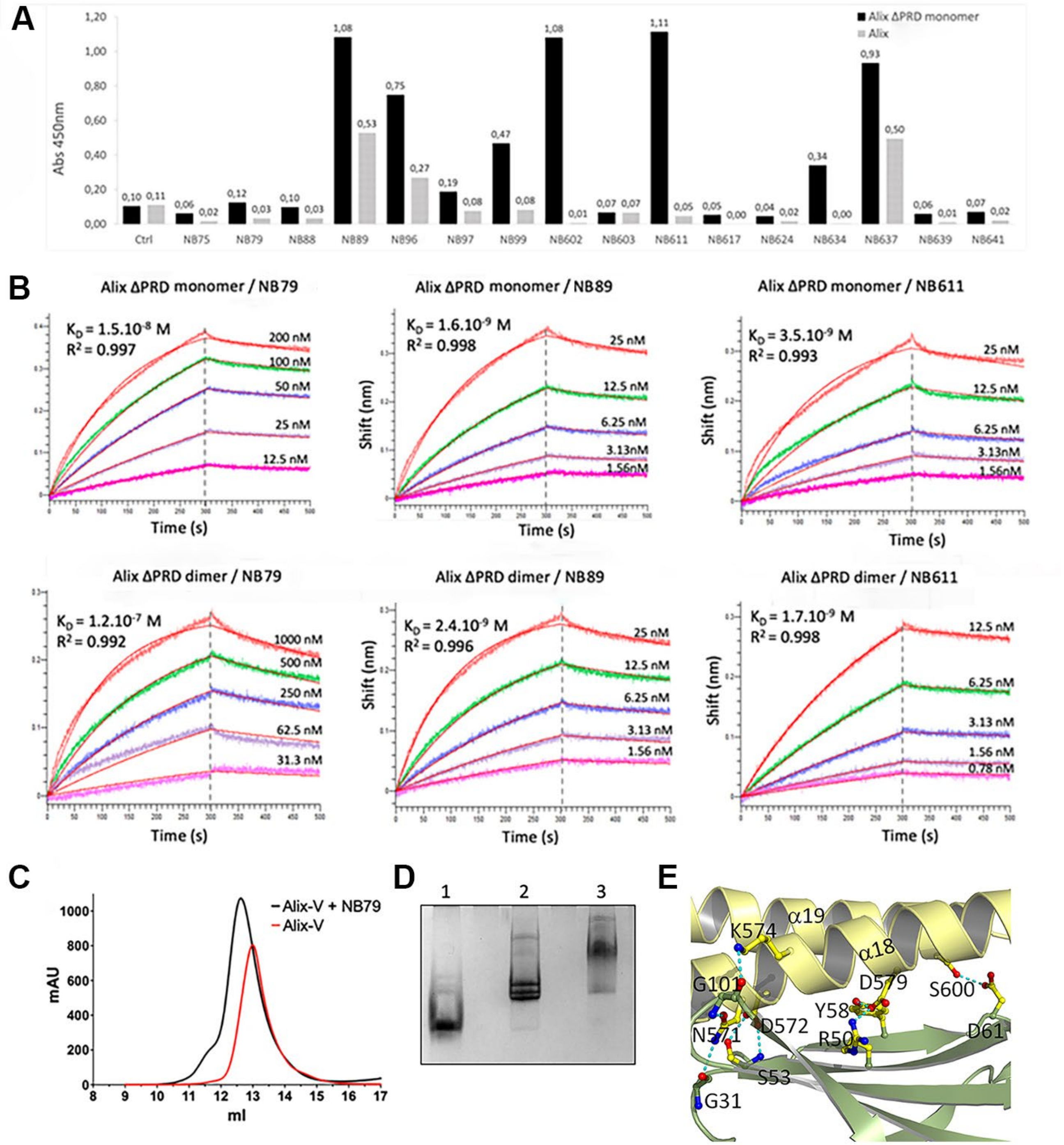
A
Alix ΔPRD monomer
Alix
Abs 450nm
Ctrl
NB75
NB79
NB88
NB89
NB96
NB97
NB99
NB602
NB603
NB611
NB617
NB624
NB634
NB637
NB639
NB641
B
Alix ΔPRD monomer / NB79
$K_D$ = 1.5.10$^{-8}$ M
$R^2$ = 0.997
Alix ΔPRD monomer / NB89
$K_D$ = 1.6.10$^{-9}$ M
$R^2$ = 0.998
Alix ΔPRD monomer / NB611
$K_D$ = 3.5.10$^{-9}$ M
$R^2$ = 0.993
Alix ΔPRD dimer / NB79
$K_D$ = 1.2.10$^{-7}$ M
$R^2$ = 0.992
Alix ΔPRD dimer / NB89
$K_D$ = 2.4.10$^{-9}$ M
$R^2$ = 0.996
Alix ΔPRD dimer / NB611
$K_D$ = 1.7.10$^{-9}$ M
$R^2$ = 0.998
Shift (nm)
Time (s)
C
Alix-V + NB79
Alix-V
mAU
ml
D
1
2
3
E
K574
α19
α18
G101
D579
S600
Y58
N571
D572
R50
D61
S53
G31

**Figure S2. Characterization of Alix-V-domain-specific llama nanobodies**.

**(A)** ELISA analyses showing interaction of 16 nanobodies as indicated with AlixΔPRD (monomer) and Alix (full length monomer); Ctr, negative control.

**(B)** Bio-layer interferometry (BLI) analyses of NB79, NB89 and NB611 binding to AlixΔPRD monomers and AlixΔPRD dimers. Binding was fit to a 1:1 model employing 5 concentrations of NB79, NB89 and NB611 as indicated. Experiments were performed in triplicates, n=3, mean ± SD. $K_a$, $k_d$ and calculated $K_D$s are reported in Table S1.

**(C)** Complex formation of NB79 and dimeric Alix-V. Size exclusion chromatography (SEC) of Alix-V and Alix-V-NB79 complexes on a Superdex 200 Increase column reveals a size increase (shift) of the central peak upon complex formation.

**(D)** Native gel electrophoresis shows distinct bands for Alix-V monomers (lane 1) and dimers (lane 2) as well as a band shift of the Alix-V dimer upon complex formation with NB79 (lane 3).

**(E)** Ribbon diagram of a close-up of the NB79 (green) interaction with α18 of dimeric Alix-V (yellow). Interacting residues are shown as sticks and hydrogen bonds are indicated by dashed lines (left panel).

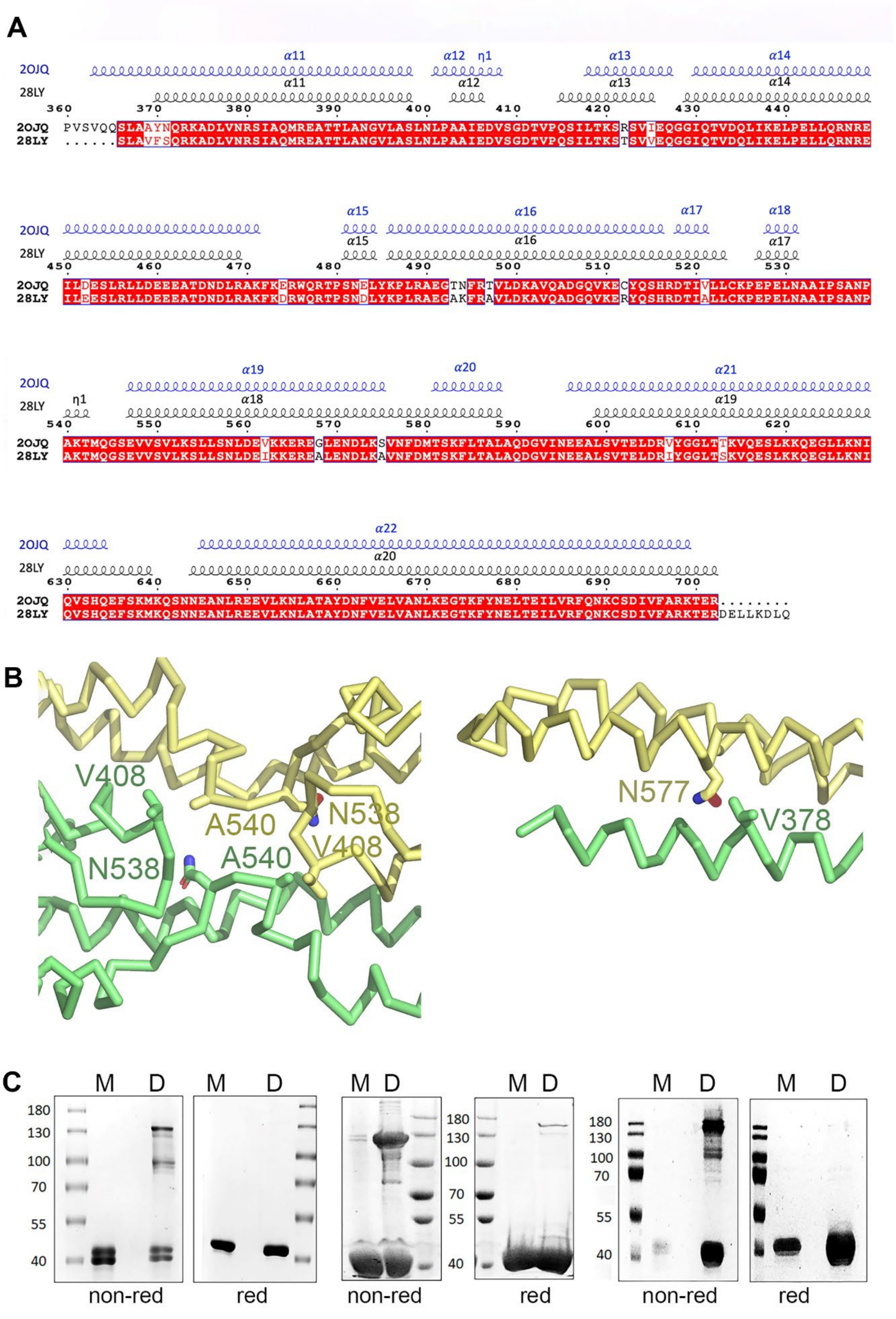
A
2OJQ
28LY
α11
α12
η1
α13
α14
2OJQ PVSVQQSLAAYNQRKADLVNRSIAQMREATTLANGVLASLNLPAAIEDVSGDTVPQSILTKSRSVTEQGGIQTVDQLIKELPELLQRNRE
28LY ......SLAVFSQRKADLVNRSIAQMREATTLANGVLASLNLPAAIEDVSGDTVPQSILTKSTSVVEQGGIQTVDQLIKELPELLQRNRE
α15
α16
α17
α18
2OJQ ILDESLRLLDEEEATDNDLRAKFKERWQRTPSNELYKPLRAEGTNFRTVLDKAVQADGQVKECYQSHRDTIVLLCKPEPELNAAIPSANP
28LY ILEESLRLLDEEEATDNDLRAKFKDRWQRTPSNDLYKPLRAEGAKFRAVLDKAVQADGQVKERYQSHRDTIALLCKPEPELNAAIPSANP
α19
α20
α21
α18
α19
2OJQ AKTMQGSEVVSVLKSLLSNLDEVKKEREGLENDLKSVNFDMTSKFLTALAQDGVINEEALSVTELDRVYGGLTTKVQESLKKQEGLLKNI
28LY AKTMQGSEVVSVLKSLLSNLDEIKKEREALENDLKAVNFDMTSKFLTALAQDGVINEEALSVTELDRIYGGLTSKVQESLKKQEGLLKNI
α22
α20
2OJQ QVSHQEFSKMKQSNNEANLREEVLKNLATAYDNFVELVANLKEGTKFYNELTEILVRFQNKCSDIVFARKTER........
28LY QVSHQEFSKMKQSNNEANLREEVLKNLATAYDNFVELVANLKEGTKFYNELTEILVRFQNKCSDIVFARKTERDELLKDLQ
B
V408
A540
N538
V408
N538
A540
N577
V378
C
M D M D M D M D M D M D
180 130 100 70 55 40
non-red
red
non-red
red
non-red
red

**Figure S3. Comparison of the V-domain monomer and dimer and dimer structure validation.**
**(A)** Comparison of the secondary structure elements of the V-domain from monomeric (pdb 2OJQ) and dimeric (pdb 28LY) Alix-V structures.
**(B)** Mutations of residues into cysteines positioned to produce disulfide-linked dimers. Combinations of cysteine pairs of hinge region residues N538, V408 and A540 (right panel) positions their SH groups at distances of 2.58 Å (V408C-A540C) and between 3.4 to 4.1 Å (N538C-A540C). Cysteine mutations of residues V378C and N577C positions their respective SH groups at disulfide bonding distance (2.35 Å).
**(C)** (left panel) SDS-PAGE analyses of monomeric (M) and dimeric (D) Alix-$V_{V408C\text{-}A540C}$ shows the formation of disulfide-linked dimers under non-reducing conditions for the dimeric form but not for the monomeric form; both monomer and dimer reveal one band corresponding to the size of the monomer under reducing SDS-PAGE conditions.
(middle panel) SDS-PAGE analyses of Alix-$V_{N538C\text{-}A540C}$ shows the formation of disulfide-linked dimers under non-reducing conditions for the dimeric form; both monomer and dimer reveal one band corresponding to the size of the monomer under reducing SDS-PAGE conditions.
(right panel) SDS-PAGE analyses of Alix-$V_{V378C\text{-}N577C}$ shows the formation of disulfide-linked dimers under non-reducing conditions for the dimeric form; both monomer and dimer reveal one band corresponding to the size of the monomer under reducing SDS-PAGE conditions.

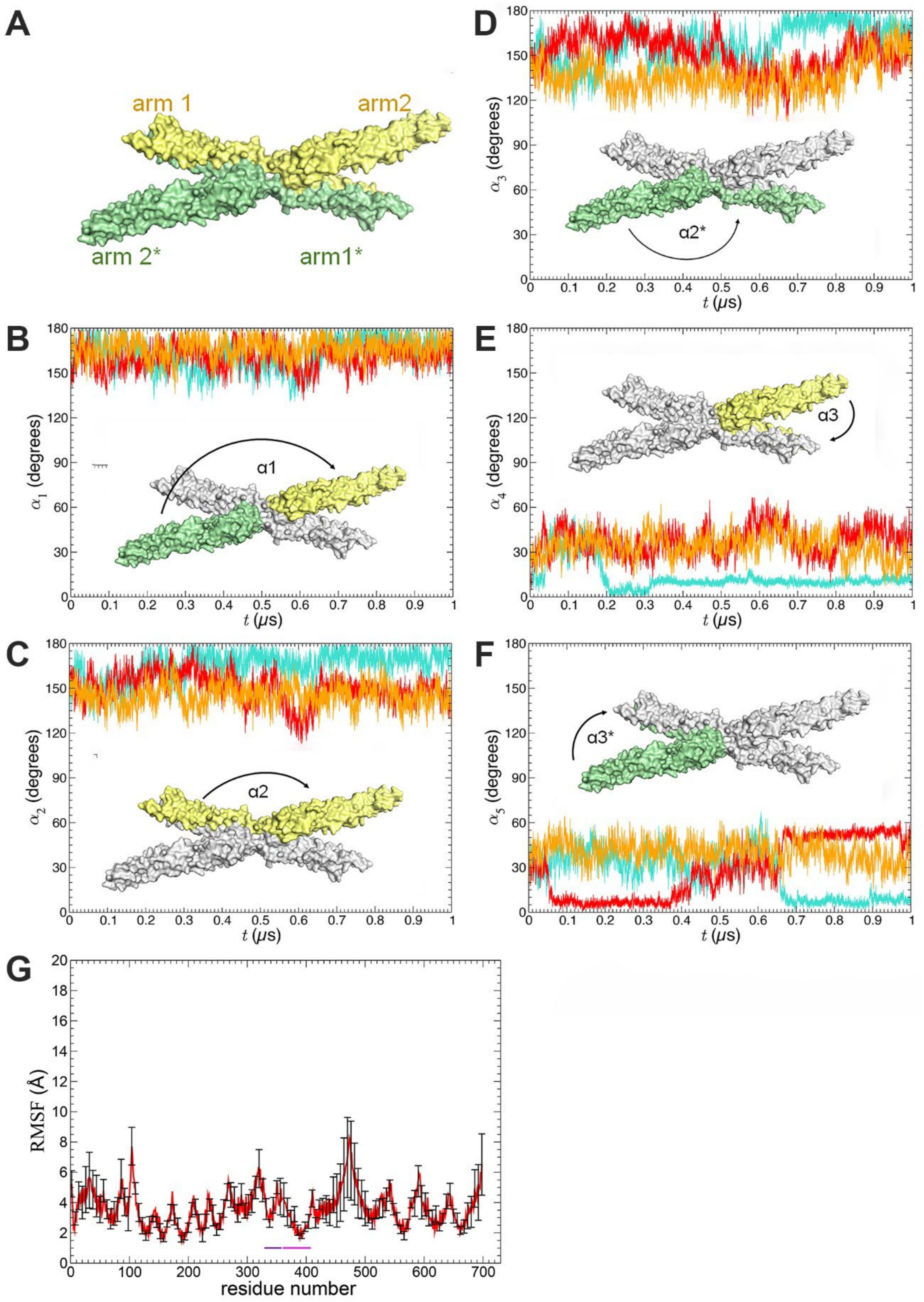
A
arm 1
arm2
arm 2*
arm1*
B
α1
$\alpha_1$ (degrees)
$t$ (μs)
C
α2
$\alpha_2$ (degrees)
$t$ (μs)
D
α2*
$\alpha_3$ (degrees)
$t$ (μs)
E
α3
$\alpha_4$ (degrees)
$t$ (μs)
F
α3*
$\alpha_5$ (degrees)
$t$ (μs)
G
RMSF (Å)
residue number

**Figure S4. Molecular dynamics simulation reveals the conformational flexibility of dimeric Alix-V.**

**(A)** Space filling representation of dimeric Alix showing arms 1 and 2 and its symmetry related arm 1* and arm 2*. The α1, α2 and α3 angles and their symmetry mates (*) are indicated.

The following MD simulation plots show the degree of fluctuations during 1 μs between the indicated angles α (°)

**(B)** α1 of arm 2 and arm 2*

**(C)** α2 of arm 1 and arm 2 and **(D)** its symmetry related angles α2*

**(E)** α3 of arm 1 and arm 2* and **(F)** its symmetry related angles α3*

**(G)** Plot of the per residue root mean square fluctuation (RMSF) (Å) of the Alix-Bro-V structure (AlixΔPRD) (pdb 2OEV) demonstrating local flexibility. Plotted values are averages from five independent simulations.

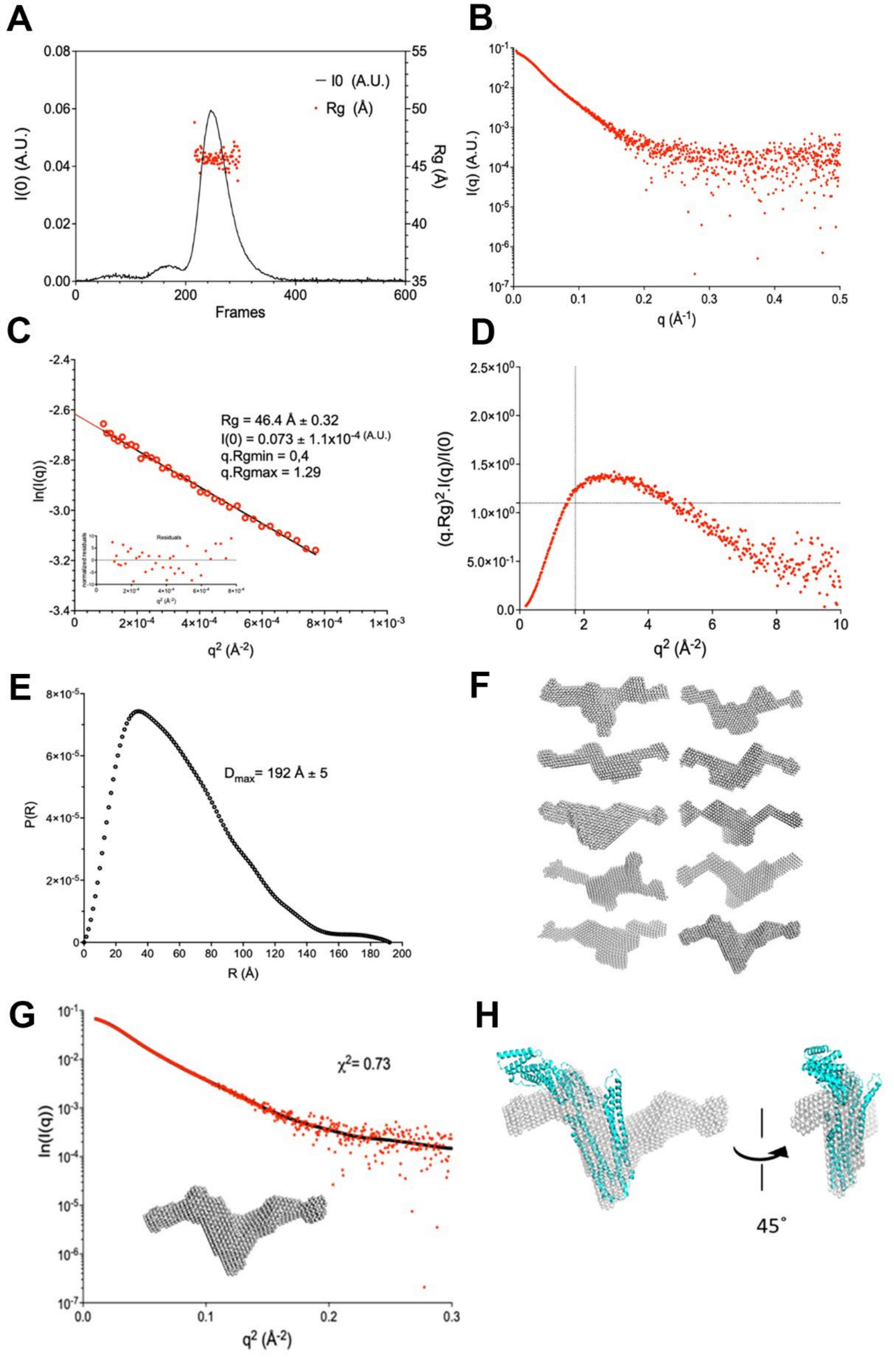

A
I0 (A.U.)
Rg (Å)
I(0) (A.U.)
Rg (Å)
Frames
B
I(q) (A.U.)
q (Å-1)
C
Rg = 46.4 Å ± 0.32
I(0) = 0.073 ± 1.1x10-4 (A.U.)
q.Rgmin = 0,4
q.Rgmax = 1.29
ln(I(q))
Residuals
q2 (Å-2)
D
(q.Rg)2.I(q)/I(0)
q2 (Å-2)
E
Dmax= 192 Å ± 5
P(R)
R (Å)
F
G
χ2= 0.73
ln(I(q))
q2 (Å-2)
H
45˚

**Figure S5. SAXS analyses of full-length Alix.**

**(A)** SEC-MALLS analysis of Alix-V reveals the Rg of monodisperse Alix.

**(B)** Small angle X-ray scattering data of Alix.

**(C)** Guinier plot analysis, revealing an Rg of 46.4 Å.

**(D)** The dimensionless Kratky plot displays a defined maximum and decays at high q, indicating a predominantly folded and compact particle with some flexibility, consistent with a flexible/disordered PRD.

**(E)** P(r) functions of full-length Alix shows a Dmax of 192 Å.

**(F)** Ab initio calculated model of full-length ALIX reveals an ensemble of similar models.

**(G)** The model with the best fit, lowest $chi^2$ shows an extended V-shaped envelope.

**(H)** The structure of Alix-Bro-V (pdb 2EOV) was placed into the envelope demonstrating that the Bro1 domain and arm 1 of the V-domain could fit into the left arm of the extended V-shaped envelope by rotation and the right extension of the envelope could accommodate arm 2 of the V-domain plus the PRD extension.

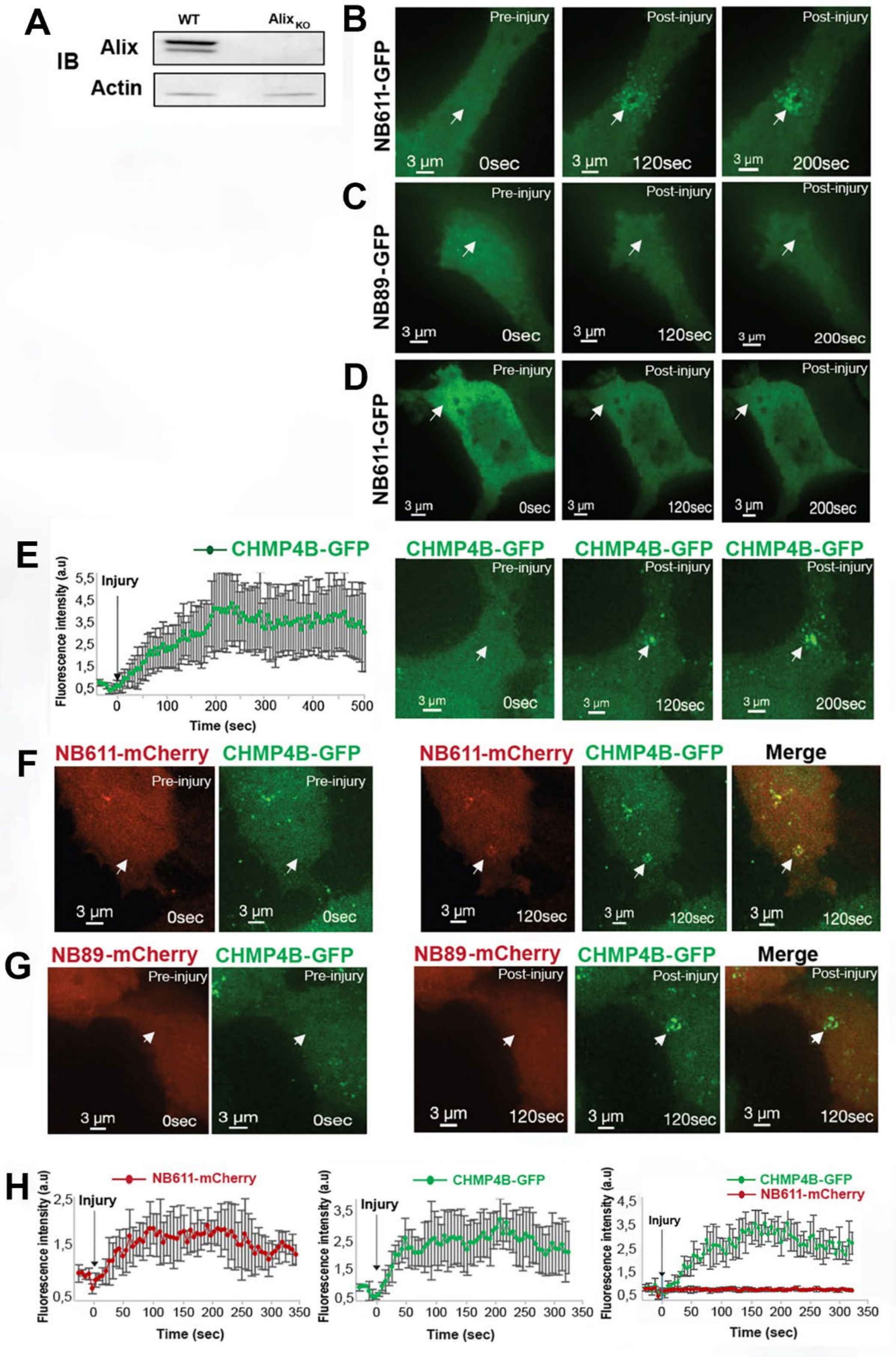
A
WT
Alix KO
IB
Alix
Actin
B
NB611-GFP
Pre-injury
Post-injury
3 µm
0sec
120sec
200sec
C
NB89-GFP
D
NB611-GFP
E
CHMP4B-GFP
Injury
Fluorescence intensity (a.u)
Time (sec)
F
NB611-mCherry
CHMP4B-GFP
Merge
G
NB89-mCherry
H
NB611-mCherry
CHMP4B-GFP

**Figure S6. NB611-GFP recognizes endogenous Alix at membrane repair sites, which in turn recruits CHMP4B.**

**(A)** Western blot analyses of Alix expression in HEK293 WT and Alix$_{_KO}$ cells. Left lane, endogenous Alix expression in HEK293 cells; right lane, endogenous Alix expression in HEK293 Alix$_{_KO}$ cells. Lower panel, both cells express actin.

**(B)** HEK293 cells were transfected with NB611-GFP, **(C)** NB89-GFP and **(D)** HEK293 Alix$_{_KO}$ cells were transfected with NB611-GFP. Plasma membrane of transfected cells was injured using a pulsed laser and cells were imaged by confocal microscopy. This revealed recruitment of NB611-GFP in HEK293 cells at 120 sec (middle panel) and 200 sec (right panel) post injury **(B)** and **(C)** no recruitment of NB89-GFP post injury at 120 sec and 200 sec and **(D)** no recruitment of NB611-GFP in HEK293 Alix$_{_KO}$ cells upon injury at 120 sec and 200 sec. White arrows indicate the sites of injury. Scale bar is 3 µm.

**(E)** HeLa CHMP4B (CHMP4B-GFP) expressing cells (left panel) were injured using a pulsed laser. Confocal images show CHMP4B recruitment at the plasma membrane injury site at 120 sec (middle panel) and 200 sec (right image panel). The graph (right panel) shows the kinetics of CHMP4B-GFP fluorescence intensity at the injury site; injury at 0 sec is indicated by a flash.

**(F)** HeLa CHMP4B expressing cells were transfected with NB611-mCherry and the plasma membrane was injured using a pulsed laser followed by confocal microscopy imaging; left panels show pre-injury cytosolic expression of NB611-mCherry and CHMP4B-GFP; right panels demonstrate post injury NB611-mCherry staining of Alix, CHMP4B-GFP recruitment and their co-localization, indicative of endogenous Alix recruiting CHMP4B.

**(G)** HeLa CHMP4B (CHMP4B-GFP) expressing cells were transfected with NB89-mCherry and the plasma membrane was injured using a pulsed laser followed by confocal microscopy imaging; the left panels show cytosolic expression of NB611-mCherry and CHMP4B-GFP pre-injury; right panels, localization of NB89-mCherry post-injury, CHMP4B-GFP recruitment and their co-localization, indicating that NB80 cannot detect endogenous Alix recruiting CHMP4B.

**(H)** Graphs show the fluorescence intensities of NB611-mCherry (left panel, related to F), CHMP4B-GFP (middle panel, related to F) and NB89-mCherry, CHMP4B-GFP (right panel, related to G) at the injury sites; injury at 0 sec is indicated by an arrow.

Scale bars correspond to 3 µm.

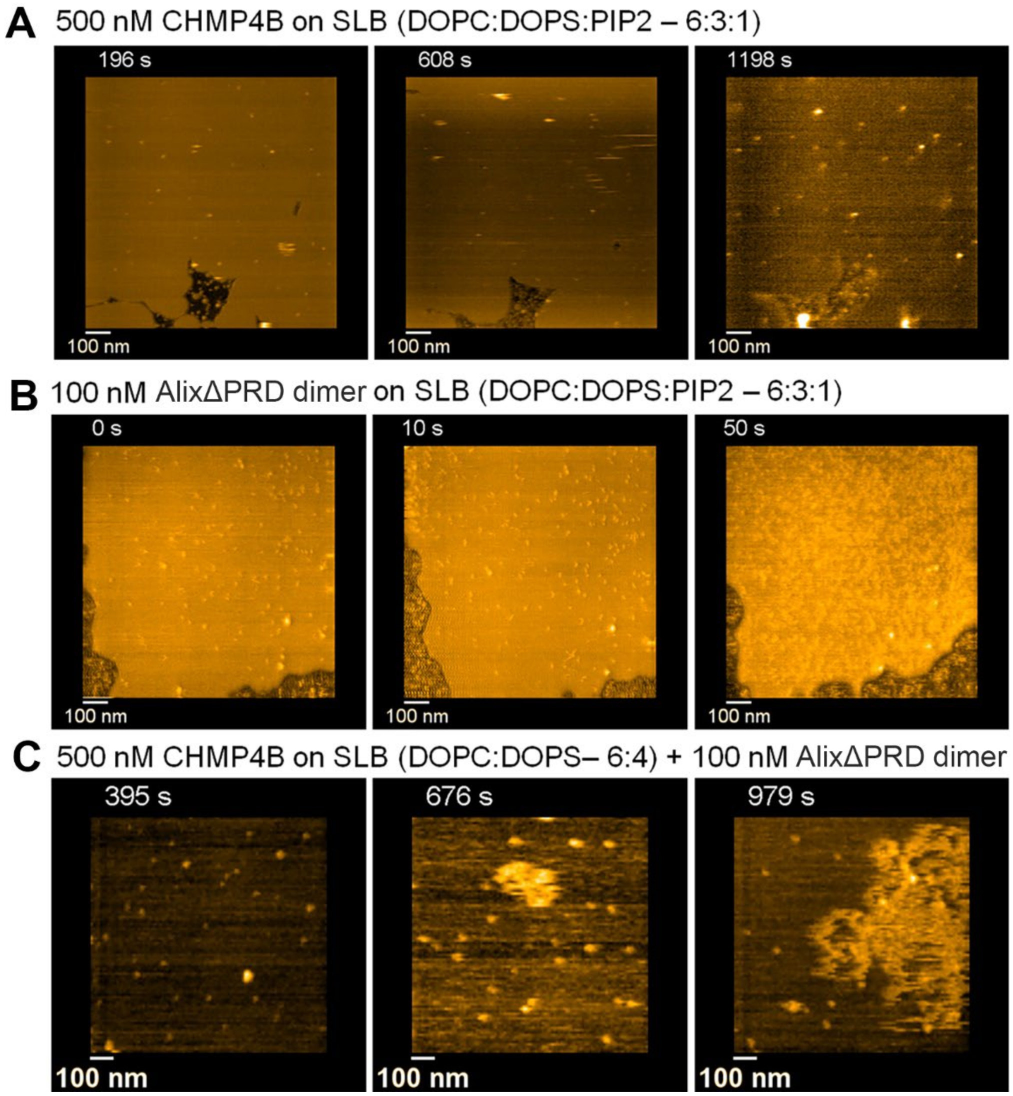


**Figure S7. HS-AFM imaging of CHMP4B and AlixΔPRD.**

**(A)** Time-lapse HS-AFM images showing that CHMP4B alone does not polymerize on supported lipid bilayers composed of DOPC:DOPS:PIP2 (6:3:1 molar ratio) at a concentration of 500 nM; related to **movie 1**.

**(B)** Time-lapse HS-AFM images show dimeric AlixΔPRD alone on supported lipid bilayers, including PIP2, do not lead to higher order structures; related to **movie 2**.

**(C)** Time-lapse HS-AFM images of 500 nM CHMP4B and AlixΔPRD dimers on supported lipid bilayers composed of DOPC:DOPS (6:4 molar ratio) (i.e. without PIP2) show unstable oligomeric states; related to **movie 5**.

**Movie legends**

**Movie 1. MCherry-Alix imaging:** HEK 293 Alix$_{_KO}$ cells were transfected with mCherry-Alix and NB611-GFP. Plasma membrane was injured at 0 sec using a pulsed laser and imaged by confocal microscopy. See corresponding Figure 3A.

**Movie 2. NB611-GFP imaging:** HEK 293 Alix$_{_KO}$ cells were transfected with mCherry-Alix and NB611-GFP. Plasma membrane was injured at 0 sec using a pulsed laser and imaged by confocal microscopy. See corresponding Figure 3A.

**Movie 3. MCherry-Alix imaging:** HEK 293 Alix$_{_KO}$ cells were transfected with mCherry-Alix and NB89-GFP. Plasma membrane was injured at 0 sec using a pulsed laser and imaged by confocal microscopy. See corresponding Figure 3B.

**Movie 4. NB89-GFP imaging**: HEK 293 Alix$_{_KO}$ cells were transfected with mCherry-Alix and NB89-GFP. Plasma membrane was injured at 0 sec using a pulsed laser and imaged by confocal microscopy. See corresponding Figure 3B.

**Movie 5. CHMP4B-GFP imaging:** HEK293 cells were transfected with CHMP4B-GFP and mCherry Alix. Plasma membrane was injured using a pulsed laser at time 50 sec and cells were imaged using confocal microscopy. See corresponding Figure 4A.

**Movie 6. MCherry-Alix imaging:** HEK293 cells were transfected with CHMP4B-GFP and mCherry Alix. Plasma membrane was injured using a pulsed laser at time 50 sec and cells were imaged using confocal microscopy. See corresponding Figure 4A.

**Movie 7. To-PRO3 uptake imaging:** HEK293 cells were transfected with CHMP4B-GFP and mCherry Alix. Plasma membrane was injured using a pulsed laser at time 50 sec and cells were imaged using confocal microscopy. See corresponding Figure 4A.

**Movie 8. CHMP4B-GFP imaging:** HEK293 cells were transfected with CHMP4B-GFP and mCherry Alix_mut1. Plasma membrane was injured using a pulsed laser at time 50 sec and cells were imaged using confocal microscopy. See corresponding Figure 4B.

**Movie 9. MCherry-Alix_mut1 imaging**: HEK293 cells were transfected with CHMP4B-GFP and mCherry Alix_mut1. Plasma membrane was injured using a pulsed laser at time 50 sec and cells were imaged using confocal microscopy. See corresponding Figure 4B.

**Movie 10. To-PRO3 uptake imaging:** HEK293 cells were transfected with CHMP4B-GFP and mCherry Alix_mut1. Plasma membrane was injured using a pulsed laser at time 50 sec and cells were imaged using confocal microscopy. See corresponding Figure 4B.

**Movie 11. HS-AFM of CHMP4B.** HS-AFM movie of supported lipid bilayers composed of DOPC:DOPS:PIP2 (6:3:1 molar ratio) in the presence of 500 nM CHMP4B. Imaging time 2 secs per frame. See corresponding Figure 6A

**Movie 12. HS-AFM of dimeric Alix**: HS-AFM movie of supported lipid bilayers composed of DOPC:DOPS:PIP2 (6:3:1 molar ratio) in the presence of 100 nM AlixΔPRD dimers. Imaging time 2 secs per frame. Related to Figure 6.

**Movie 13. HS-AFM of monomeric Alix and CHMP4B**: HS-AFM movie of supported lipid bilayers composed of DOPC:DOPS:PIP2 (6:3:1 molar ratio) in the presence of CHMP4B and 100 nM monomeric AlixΔPRD. Imaging time 3 secs per frame. Related to Figure 6.

**Movie 14. HS-AFM of dimeric Alix and CHMP4B**: HS-AFM movie capturing the oligomerization of stable CHMP4B filaments induced by dimeric AlixΔPRD on supported lipid bilayers composed of DOPC:DOPS:PIP2 (6:3:1 molar ratio) in the presence of a mixture of 500 nM CHMP4B and 100 nM dimeric AlixΔPRD. Imaging time 2 secs per frame. See corresponding Figure 4C.

**Movie 15. HS-AFM of dimeric Alix and CHMP4B, no PIP2**: HS-AFM movie capturing the oligomerization of unstable CHMP4B filaments induced by dimeric AlixΔPRD on supported lipid bilayers composed of DOPC:DOPS (6:4, molar ratio, without PIP2) in the presence of a mixture of 500 nM CHMP4B and 100 nM AlixΔPRD dimers. Imaging time 1 sec per frame. See corresponding Figure S7.
